# Aberrant axonal pathfinding and exuberant myelination in an inducible model of neocortical heterotopia

**DOI:** 10.1101/2020.03.11.987388

**Authors:** Alice M. Li, Robert A. Hill, Jaime Grutzendler

## Abstract

Neocortical heterotopia consist of ectopic neuronal clusters that are frequently found in individuals with cognitive disability and epilepsy. However, their pathogenesis remains poorly understood due in part to a lack of tractable animal models. We have developed an inducible model of focal heterotopia that enables their precise spatiotemporal control and high-resolution optical imaging in live mice. Here we report that heterotopia are associated with striking patterns of hypermyelinated and circumferentially projecting axons around neuronal clusters. Despite their aberrant axonal patterns, *in vivo* calcium imaging revealed that heterotopic neurons remain functionally connected to other brain regions, highlighting their potential to influence global neural networks. These aberrant patterns only form when heterotopia are induced during a critical embryonic temporal window, but not in early postnatal development. Our model provides a new way to investigate heterotopia formation *in vivo* and revealed features suggesting the existence of developmentally-modulated, neuron-derived axon guidance and myelination factors.

## INTRODUCTION

Up to one third of routine postmortem examinations reveal the presence of neocortical heterotopia^1–4^, a heterogeneous group of focal cortical lamination defects characterized by abnormally positioned clusters of neurons^5,6^. Heterotopia have been linked to many neurological conditions including epilepsy, intellectual disability, and dyslexia^3,7–12^. However, comprehensive exploration of their pathogenesis and pathophysiology has been limited by a lack of tools for their investigation in the live animal. Several genetic, traumatic, chemotoxic and other neocortical heterotopia models have been described^13–21^. However, these previous models have not been utilized for cellular intravital optical imaging analyses largely due to the lack of control over the position and timing of heterotopia induction and/or limited means for targeted cell labelling.

Here, we developed a methodology enabling the specific induction and visualization of focal heterotopia in the live mammalian neocortex. Our model uses *in utero* electroporation^22–24^ combined with *in vivo* optical imaging, to generate and track layer I cortical heterotopia. When visualized using label free myelin imaging^25–27^, we identify striking patterns of aberrantly projecting and hypermyelinated axons surrounding the heterotopic neurons. These distinct patterns emerge only when the heterotopia are induced during a critical embryonic period, suggesting the presence of locally-derived, developmentally-modulated signals that initiate the abnormal structural organization and myelination of the heterotopic axons. Identification and characterization of the neuronal and glial subtypes within the induced heterotopia revealed many consistent features that are used to define spontaneously occurring heterotopia in both humans^28,29^ and rodent models^18,30–33^. Finally, using genetically encoded calcium biosensors, we reveal that heterotopic neurons display similar calcium transient frequencies as neighboring layer II/III neurons and respond to sensorimotor stimulation in behaving mice, opening new possibilities for the exploration of their influence on cortical function.

## RESULTS

### Spatiotemporally precise layer I heterotopia generation and visualization in the live mouse brain

*In utero* electroporation (IUE) is a powerful technique that permits labeling and genetic manipulation of targeted cortical neurons^22–24^. As part of the IUE procedure, a thin glass microcapillary needle is used to deliver genetic material into the lateral ventricle of embryonic stage animals for the electroporation of neuronal progenitors (Fig. 1a). While implementing IUE to label and image cortical axons *in vivo*, we unexpectedly found that cortical sites that had been injected during the procedure developed marked accumulations of labelled neuronal cell bodies in layer I of adult mice (Fig. 1c, top row). Electroporated pyramidal cells are normally found in layers II/III of cortical regions, thus their distinct presence at injected layer I sites stood out in contrast to non-injected surrounding cortical areas (Fig. 1c, top row). *In vivo* labeling of neurons with a fluorescent dye called NeuO^34^ further revealed that the IUE-labelled neuronal cell bodies constituted only a small fraction of the total neuronal cell bodies ectopically positioned (Fig.1b, c; for detailed statistics see Supplementary File 1; mean diameter of NeuO^+^ ectopic cell clusters 474 ± 174 µm s.d. in *n* = 23 mice). Abnormal superficial clustering of neuronal cell bodies is a defining feature of layer I heterotopia^1,4,28^, a subtype of neocortical heterotopia that has been linked to learning impairments in humans^7,35,36^. Their presence demonstrates that direct injections into the cortex performed as part the IUE procedure can induce molecularly tractable neocortical heterotopia *in vivo*.

**Fig. 1.**
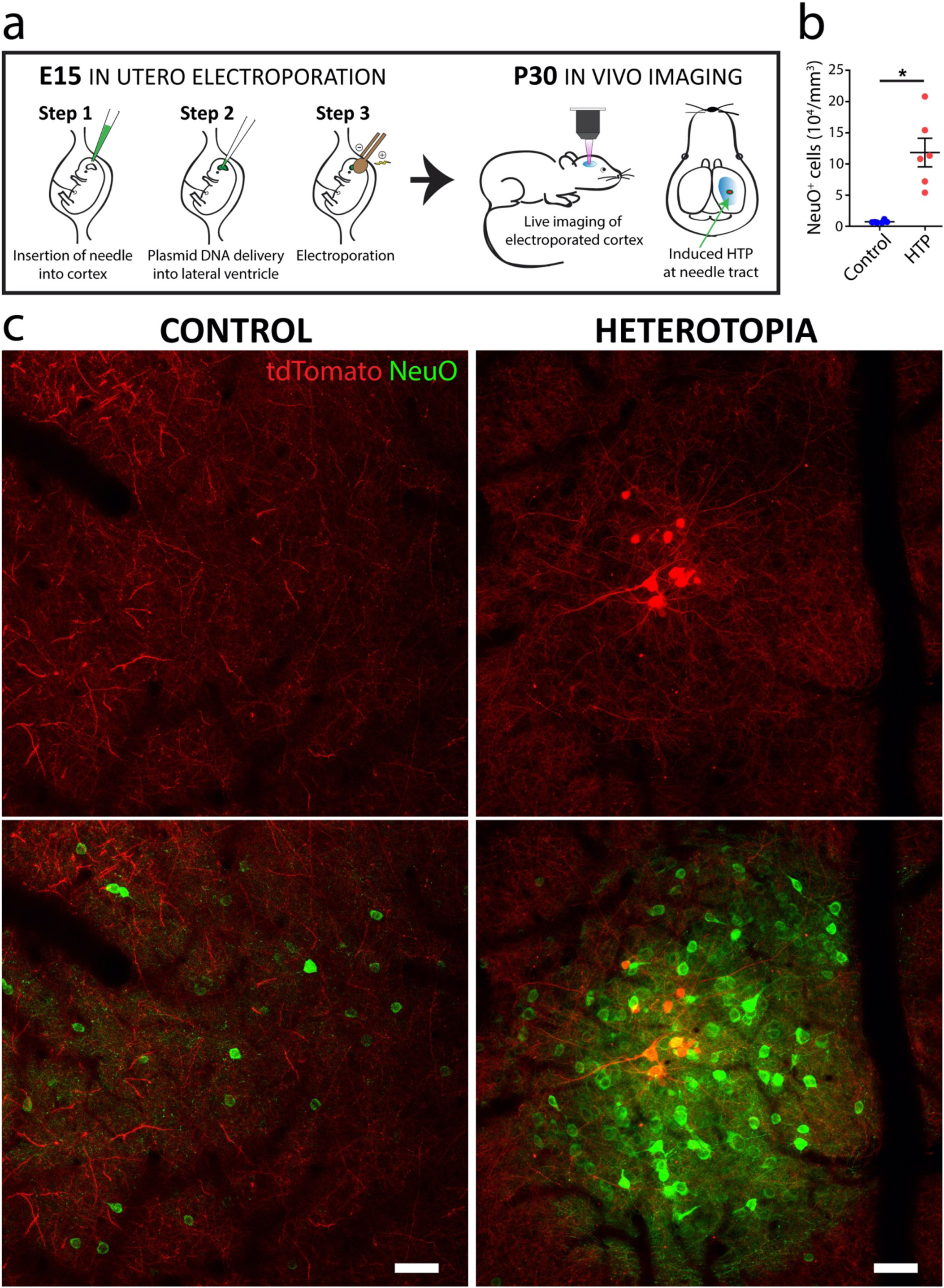
Targeted induction and high-resolution imaging of layer I heterotopia in the live mouse. **a**, Diagram showing a single neocortical heterotopion induced at the needle tract during *in utero* electroporation (IUE) at embryonic day (E15). The site is relocated postnatally for detailed investigation of the resulting layer I heterotopion by intravital imaging. **b**, Quantification showing significantly increased neuron density (*n* = 6 mice) within heterotopia compared to ipsilateral neighboring layer I control regions (Wilcoxon non-parametric, matched-pairs signed rank test *P < 0.05). Each point represents the heterotopion or control region from one animal. Horizontal lines and error bars denote mean and s.e.m, respectively. Descriptive statistics are indicated in Supplementary File 1. **c**, *In vivo* fluorescence images of a representative layer I heterotopion (right) and neighboring ipsilateral, non-injected control region in a P30 mouse (left). Notice the tightly packed cluster of fluorescently labelled neuronal cell bodies (NeuO; green). Neurons within the heterotopion can also be readily labeled during the electroporation procedure and visualized *in vivo* (tdTomato; red). In contrast, neurons in more sparsely populated control layer I regions never demonstrate IUE-mediated labelling (left column). HTP, heterotopia. Scale bars, 50 µm (**c**).

### Aberrant axonal and myelin patterns characterize layer I heterotopia formed during a critical embryonic time window

Using this methodology, we set out to examine the cellular composition of the layer I heterotopia. Interestingly, we observed markedly aberrant patterns of myelinated axons exclusively associated with the ectopic neuronal clusters (Fig. 2a and Fig. 3a), as visualized *in vivo* by spectral confocal reflectance microscopy (SCoRe), a technique that enables high resolution label-free imaging of myelinated axons^25,26^. This was further corroborated by fixed tissue immunohistochemistry of myelin markers, showing increased oligodendrocyte and myelin densities at the sites of injection (Fig. 2b-d, Fig. 3e-g, and Figure 3–figure supplement 1; for detailed statistics see Supplementary File 1). On closer examination *in vivo*, we observed both myelinated and unmyelinated axons swirling in a nest-like fashion around heterotopic neurons (Fig. 2a, Fig. 3a-d, and Figure 3–figure supplement 1), often forming thick concentric borders that were most prominent towards the pial surface (Fig. 3e-g and Video 1). Radially-oriented, myelinated fiber bundles were also observed projecting out from underneath the heterotopia (Fig. 2b-c and Figure 2–figure supplement 1), similar to previous descriptions of spontaneously occurring layer I heterotopia^28–31^. Thus, by applying our methodology in combination with SCoRe microscopy and immunohistochemical analyses, we discovered that distinct axon guidance and myelin abnormalities occur in embryonically*-*induced layer I heterotopia.

**Fig. 2.**
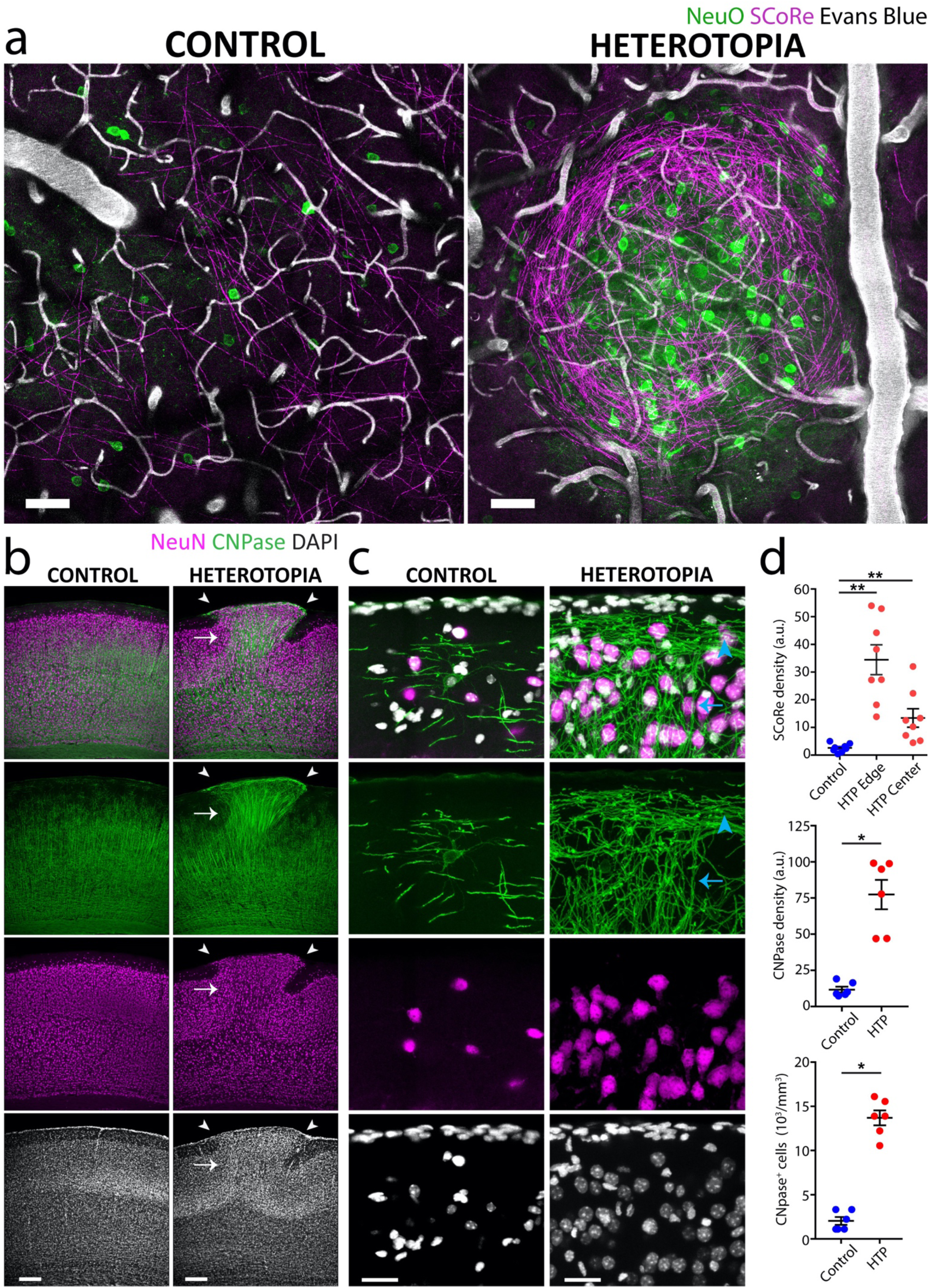
Heterotopia contain an abundance of hypermyelinated axons. **a**, *In vivo* combined fluorescence and label-free SCoRe myelin images of a representative induced heterotopion (right; same as in Figure 1c) and neighboring ipsilateral, non-injected control region (left) in a P30 mouse. The tightly packed heterotopic neuronal cell bodies (NeuO) are surrounded by a circumscribed abundance of aberrantly projecting, myelinated axon segments (SCoRe). **b**,**c** Low magnification (**b**) and high magnification (**c**) immunostainings of an induced heterotopion and its corresponding contralateral control region taken from a P30 mouse, showing increased oligodendrocyte CNPase expression associated with the heterotopion, as defined by the ectopically positioned NeuN^+^ neuronal cell bodies in layer I (**b**, white arrowheads). Notice the more horizontal orientation of myelin segments located closer to the pial surface of the heterotopion (**c**, blue arrowhead). Deeper myelin segments, in contrast, are organized in a more radial fashion (**c**, blue arrow) and fasciculate into densely myelinated fiber bundles that project through lower cortical layers (**b**, white arrow). **d**, Quantifications of SCoRe *in vivo* (*n* = 8 mice; top) and CNPase in fixed tissue (*n* = 6 mice; middle and bottom), showing increased myelination and oligodendrocyte cell body densities in induced layer I heterotopia compared to non-injected contralateral control regions. Each point represents the layer I heterotopion or control region from one animal. Horizontal lines and error bars denote mean and s.e.m, respectively (Wilcoxon matched-pairs signed rank non-parametric test; *P < 0.05, **P < 0.01). HTP, Heterotopia. Descriptive statistics are given in Supplementary File 1. Scale bars, 50 µm (**a**), 200 µm in (**b**), and 50 µm (**c**).

**Fig. 3.**
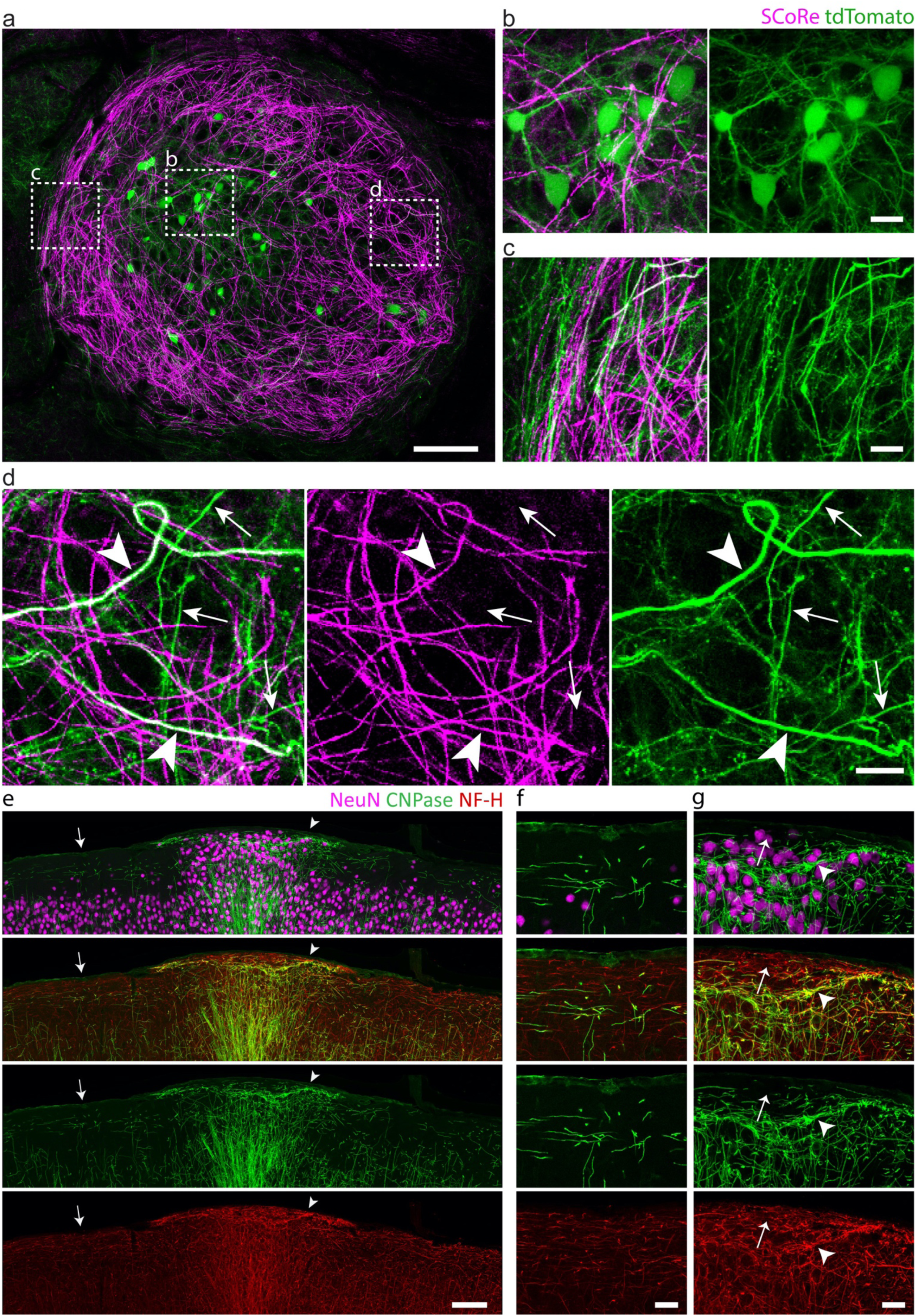
Myelinated and unmyelinated axons follow aberrantly looping and concentric paths. **a-d**, Intravital imaging of a nest-like heterotopion at P30 in (**a**) using SCoRe and confocal fluorescence microscopy showing **b**, labelled neuronal cell bodies (tdTomato; green) dispersed throughout the aberrantly oriented myelinated fibers (SCoRe; magenta) and **c**, winding axonal projections that occasionally fasciculate into bundles at the edges of the heterotopion. **d**, Myelinated (arrowheads) and unmyelinated axon segments (arrows) are observed within the heterotopion. **e-g**, Low magnification (**e**) and high magnification (**f**, **g**) immunostainings of an embryonically-induced heterotopion, confirming the presence of both myelinated (**g**, arrowhead) and unmyelinated (**f**, arrow) axon segments (**f** and **g** show high magnification images of areas indicated by arrow and arrowhead in **e**, respectively). NF-H, neurofilament heavy chain. Images are representative of experiments performed in at least six animals. Scale bars, 100 µm (**a**), 15 µm (**b-d**), 100 µm (**e**), and 25 µm (**f**, **g**).

To investigate whether the marked axonal and myelin changes were dependent on induction of heterotopia at specific developmental ages, we performed IUE at various time points during cortical development between embryonic day 14 (E14) to E17, and postnatal day 0 (P0). We found that whereas embryonic injections always resulted in similar nest-like patterns of densely myelinated fiber bundles around heterotopic neurons (Fig. 4, Fig. 5b), P0 injections did not result in robust axon guidance or myelination abnormalities, despite the presence of ectopic neural clusters (Fig. 5a). Although postnatally-induced heterotopia tended to be smaller and more variable in size (data not shown), the axonal and myelin abnormalities were never observed even around larger P0-induced clusters (Fig. 5a). Consistent with this, we found no significant difference in the myelin or oligodendrocyte cell body densities (Fig. 5c; for detailed statistics see Supplementary File 1) between heterotopia and contralateral control regions of P0-injected mice. Likewise, immunolabelling for neurofilament heavy chain (NF-H) did not reveal aberrant accumulations of axonal fibers (Fig. 5a). Together, our data indicate previously unrecognized differences in the cellular composition and organization of neocortical heterotopia based on the timing of their induction.

**Fig. 4.**
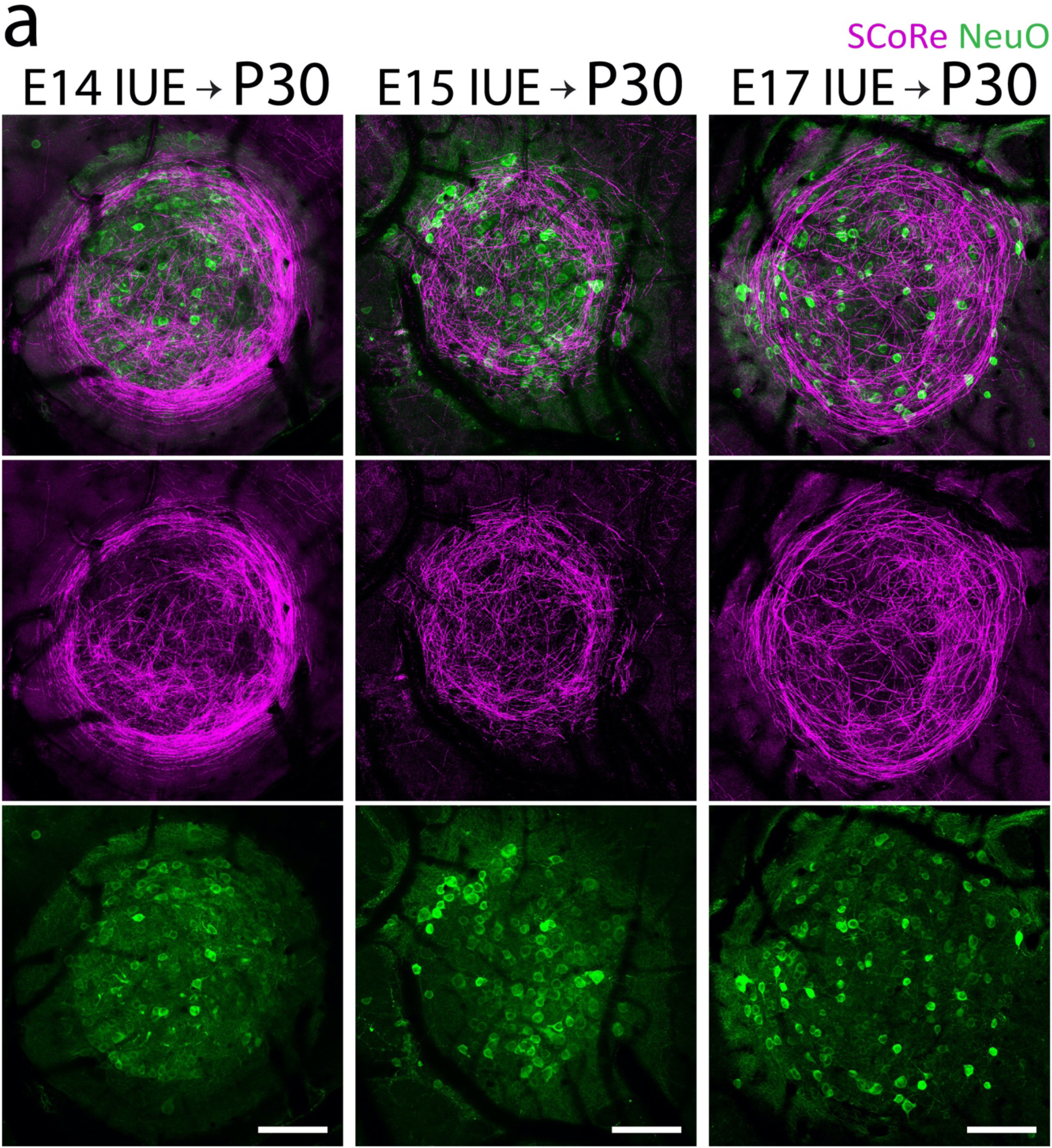
Heterotopia can be induced at various embryonic stages of corticogenesis. **a**, *In vivo* NeuO dye neuron staining (green) and SCoRe myelin imaging (magenta) of P30 mouse cortices that were previously electroporated at E14 (left column), E15 (middle column), and E17 (right column), all showing similar aberrantly projecting axons and hypermyelination encircling ectopic neuron clusters. Images are representative of observations made in at least three animals per IUE-injected age group. Scale bars, 100 µm.

**Fig. 5.**
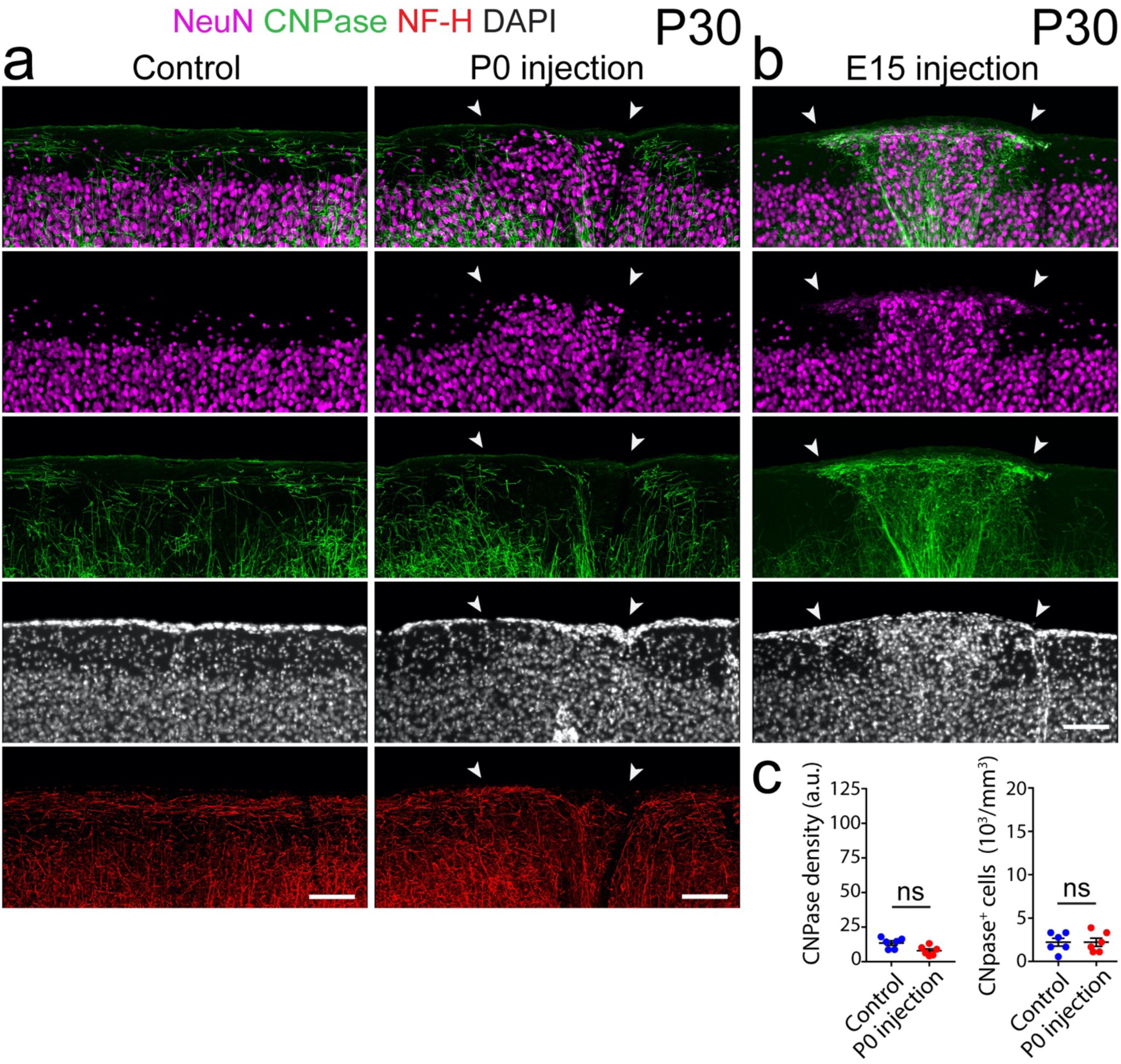
Postnatally-induced heterotopia do not develop aberrant axons or myelination. **a**, Confocal images of an immunolabelled coronal section from P30 mouse cortex, showing no changes in myelination (CNPase) or axonal pathfinding (NF-H) associated with a P0-induced heterotopion (NeuN; arrowheads), compared to its contralateral control region (left). **b**, In contrast, neuronal heterotopia induced *in utero* at E15 (NeuN; arrowheads) show dramatic changes in local layer I myelin (CNPase) expression and patterning. **c**, Quantifications of CNPase expression, showing no significant differences in the myelin or oligodendrocyte cell body densities (*n* = 6 animals) between postnatally-induced heterotopia and control regions. Each point represents the P0-induced layer I heterotopion or control region from one animal, with horizontal lines and error bars denoting mean and s.e.m, respectively. Wilcoxon matched-pairs signed rank non-parametric test was used to determine significance; NS, no significance. NF-H, neurofilament heavy chain. Descriptive statistics are given in Supplementary File 1. Scale bars, 100 µm (**a**, **b**).

### Astrocyte and microglia cell densities are not altered in induced heterotopia

Given the striking abnormalities in oligodendrocyte production, myelination and axon pathfinding observed in the embryonically-induced heterotopia, we next wondered whether other glial cell types such as astrocytes and microglia also exhibited altered morphology or density. Immunolabelled cortical brain sections against Aldh1L1 revealed no significant difference in the astrocyte cell density between layer I heterotopia and corresponding contralateral control regions (Fig. 6a-c; for detailed statistics see Supplementary File 1). Similarly, using Iba1 immunolabelling, we found no regional differences in microglia density or morphology (Fig. 6d-f; for detailed statistics see Supplementary File 1). Moreover, there was no evidence of pronounced astrocytic or microglial accumulation or altered cellular morphology at the borders of the heterotopia suggesting that a glial scar had not formed due to the embryonic injection. These data suggest that while heterotopia induce marked oligodendrocyte generation, astrocytes and microglia are not significantly influenced by potential local factors derived from ectopic neuronal clusters.

**Fig. 6.**
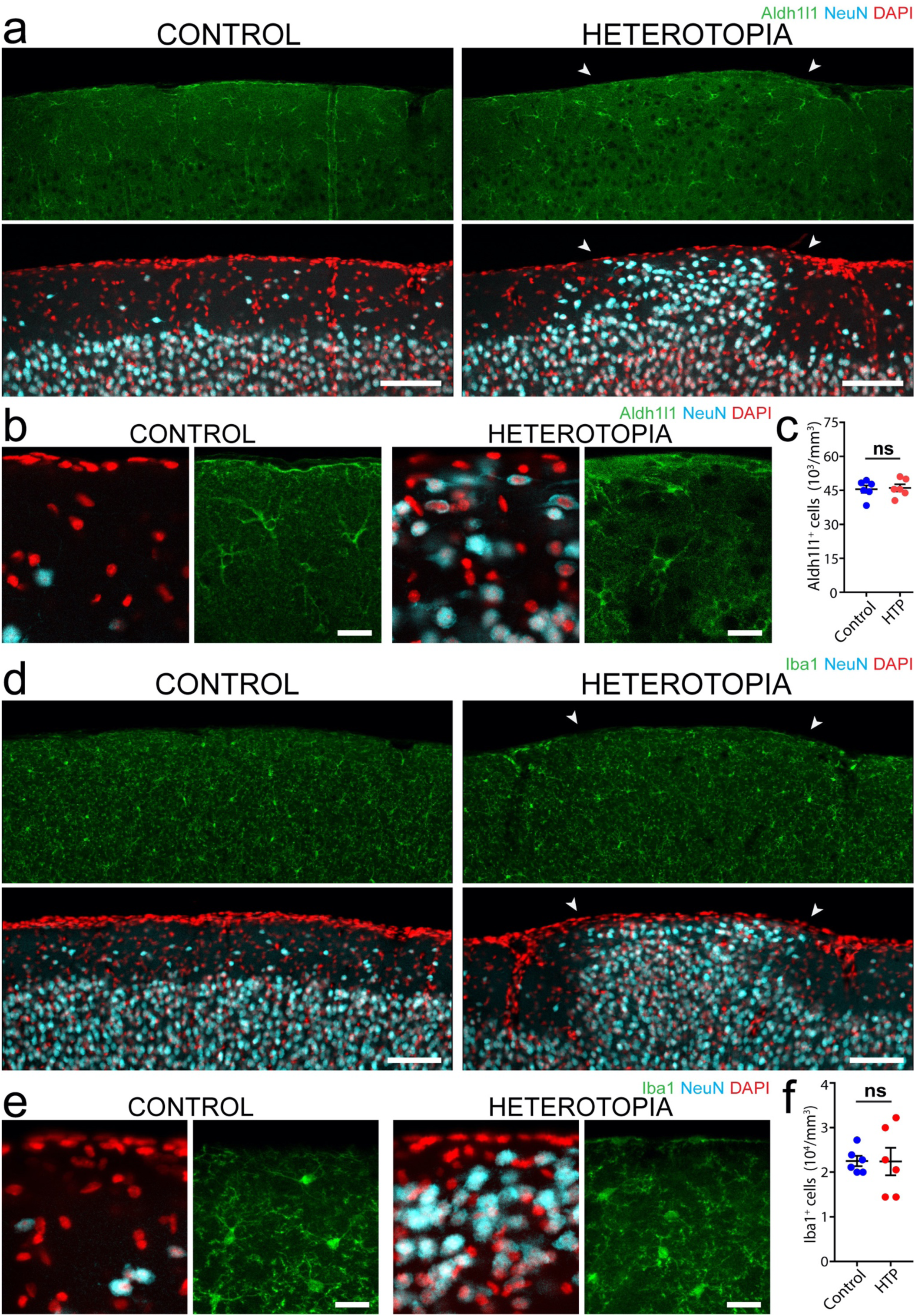
Astrocytes and microglia retain a normal density despite aberrant axonal and myelin distribution. **a-b**, Low magnification (**a**) and high magnification (**b**) images of Aldh1l1-positive astrocytes in P30 mouse cortex, showing similar cell densities in layer I heterotopia (right images, arrowheads) compared to corresponding contralateral control regions (left images). **c**, Quantifications using P30 mouse forebrain tissue showing no significant difference in astrocyte cell density (Wilcoxon non-parametric matched-pairs, signed rank test, *n* = 6 animals; ns, no significance). **d-e** Low magnification (**d**) and high magnification (**e**) images of Iba1-immunolabelled P30 mouse forebrain, showing no difference in microglia cell densities between heterotopia (right images, arrowheads) and contralateral control regions (left images). **f**, Quantifications using P30 mouse forebrain tissue demonstrating no significant difference in microglia cell density between layer I heterotopia and corresponding contralateral control regions (Wilcoxon matched-pairs, signed rank non-parametric test, *n* = 6 animals; ns, no significance). For graphs in (**c**) and (**f**), each point represents an embryonically-induced layer I heterotopion or control region from one animal, with horizontal lines and accompanying error bars denoting mean and s.e.m., respectively. Descriptive statistics are given in Supplementary File 1. HTP, heterotopia. Scale bars, 100 µm (**a**), 20 µm (**b**), 100 µm (**d**), and 20 µm (**e**).

### Heterotopia contain a variety of excitatory and inhibitory neurons born at different embryonic ages

To characterize the neural composition of the induced heterotopia, we examined the expression of neuronal subtype markers Cux1, Tle4, and GAD-67 (Fig. 7). Cux1 is a transcription factor predominantly expressed in callosal projection neurons in layers II-IV^37^, whereas Tle4 is primarily restricted to deeper corticothalamic projection neurons of layers V and VI^38,39^. We found both Cux1^+^ and Tle4^+^ neurons present to varying degrees in all layer I heterotopia (Fig. 7b, c and Figure 7–figure supplement 1), consistent with a mixed population of excitatory projection neurons from different cortical layers. Further analyses revealed the presence of GAD67^+^ interneurons, which occurred at similar densities within heterotopia as in corresponding layer I control regions (Fig. 7d-f; for detailed statistics see Supplementary File 1). In addition, layer I heterotopia always contained GAD67^+^ puncta, consistent with inhibitory synapses (Fig. 7d, e). Together, our findings suggest that induced layer I heterotopic neural clusters comprise a diverse cohort of glutamatergic and GABAergic neurons, in line with the heterogeneous population of neuronal subtypes previously described in those occurring spontaneously^31,33,40,41^.

**Fig. 7.**
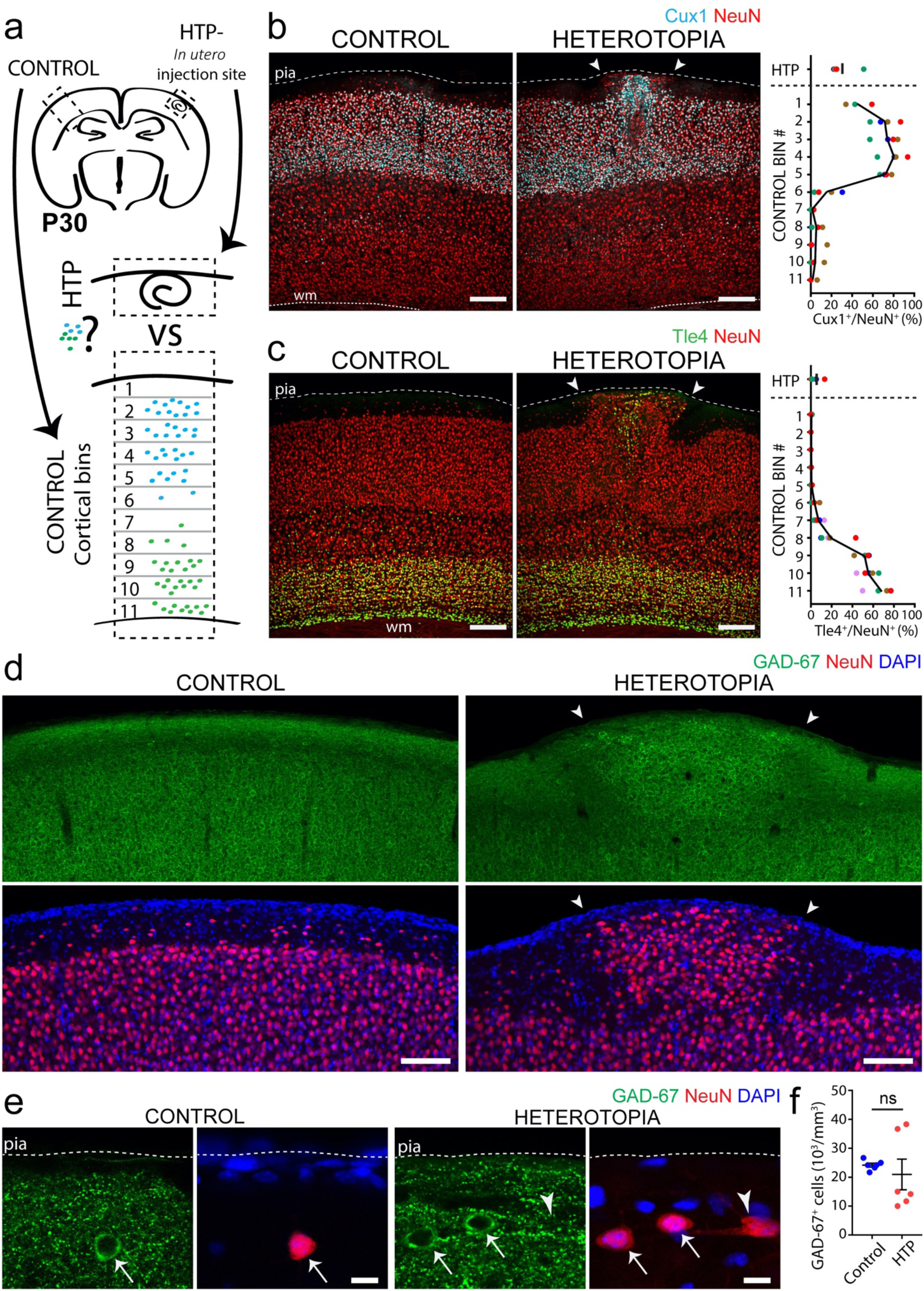
Heterotopia contain a mixed population of excitatory and inhibitory neurons. **a**, Approach for quantifying layer-specific neuronal marker expression (symbolized by blue and green dots) in layer I heterotopia and across corresponding contralateral control cortices in P30 mice. Contralateral control cortices are subdivided into 11 equally-sized bins that together span the entire thickness of the cortex. **b-c** Both Cux1^+^ (**b**, cyan) and Tle4^+^ (**c**, green) neuronal cell bodies occur within layer I heterotopia (right images, arrowheads). In contralateral control regions (left images), Cux1 is concentrated in more superficial cortical areas corresponding to layers II/III, whereas deeper areas corresponding to layers V/VI encapsulate the majority of Tle4 labelling. Quantifications (far right) show the percentages of NeuN^+^ cell bodies that also express Cux1 (top) or Tle4 (bottom) within heterotopia and across corresponding contralateral control bins. Each dot corresponds to the layer I heterotopion or control bin of a single animal, with all dots of the same color belonging to the same animal (*n* = 4 animals for Cux1; *n* = 5 animals for Tle4). The black line denotes the mean. WM, white matter. **d**,**e** Low (**d**) and high (**e**) magnification z-projections of GAD-67-immunostained P30 mouse forebrain, showing the presence of GAD-67-labelled cells and puncta in both layer I heterotopia (right images, arrowheads) and corresponding contralateral control regions (left images). **f**, Quantifications from P30 mouse tissue showing no significant difference in GAD-67^+^ neuronal cell density between layer I heterotopia and contralateral control regions (Wilcoxon matched-pairs, signed rank non-parametric test, *n* = 6 animals; ns, no significance). Each point represents the layer I heterotopion or contralateral control region from one animal, with horizontal lines and accompanying error bars denoting mean and s.e.m., respectively. HTP, heterotopia. Descriptive statistics are given in Supplementary File 1. Scale bars 200 µm (**b**, **c**), 100 µm (**d**), and 10 µm (**e**).

We next used birthdating techniques to determine whether neurons within the induced heterotopia originate at similar or different time points during development. 5-Ethynyl-2’-deoxyuridine (EdU) pulse labelling at E11.5 revealed small but distinct populations of EdU^+^NeuN^+^ cells in all embryonically-generated layer I heterotopia (Figure 7–figure supplement 2), suggesting that heterotopia contain neurons born during early corticogenesis^42,43^. We also observed cells “birthdate-labelled” via IUE^22^ at E15 (∼90% heterotopia in *n* > 30 mice; Fig. 1c) in heterotopia, suggesting the additional presence of later-born neurons. These data indicate that induced heterotopia contain neurons born at various stages of cortical development.

### Heterotopic neurons exhibit similar calcium dynamics as neighboring normotopic neurons

Despite their linkage to a wide spectrum of neurological disorders, the cellular dynamics and mechanisms by which individual heterotopic cells contribute to neural circuit dysfunction are poorly understood. We examined the calcium activity of single cortical neurons in induced heterotopia and in surrounding, non-injected cortical layer II/III regions of awake head-fixed mice using the genetically encoded calcium indicator, GCaMP6f. Surprisingly, although we found highly variable calcium spike patterns of individual cells within heterotopia (Fig. 8a-c and Video 2), there were no significant differences in their overall calcium spike event frequency, variance, or synchrony compared to surrounding non-heterotopic layer II/III regions (Fig. 8d; for detailed statistics see Supplementary File 1). Furthermore, we did not identify any epileptiform activity in our imaging sessions, although we cannot rule out that continuous recordings could have revealed sporadic aberrant activity (Fig. 8a-c and Video 2). Interestingly, we found that mice that had been startled with brief whisker stimulation consistently responded via neuronal calcium spikes within layer I heterotopia (Fig. 8e, f and Video 3). This result indicates that heterotopic neurons are connected to other brain areas and could thus influence network function.

**Fig. 8.**
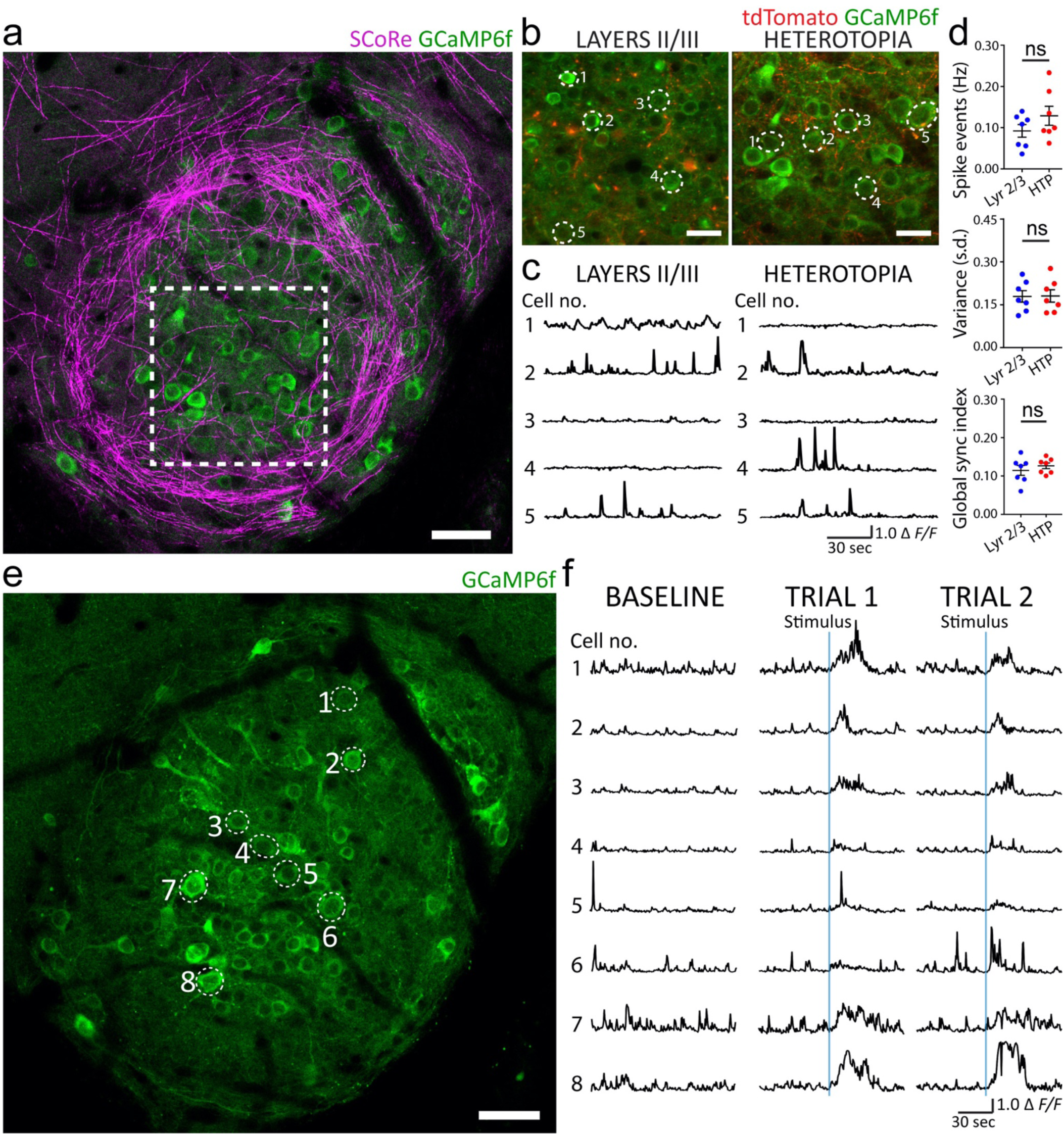
Heterotopic neurons display calcium dynamics similar to adjacent layer II/III neurons and respond to sensorimotor input. **a-c** *In vivo* time lapse imaging of layer I heterotopic neurons expressing the calcium sensor GCaMP6f in an awake, head-fixed mouse. Dotted lines in (**a**) indicate an analyzed heterotopic region that is displayed in (**b**). **c**, Example traces of spontaneous neuronal calcium transients from layer I heterotopic (right) and ipsilateral layer II/III non-heterotopic (left) regions displayed in (**b**). **d**, Quantifications showing no significant difference in calcium spike event frequency (Hz), variance (s.d.) or global synchronization index between layer I heterotopia and adjacent layer II/III non-heterotopic regions. Each dot corresponds to the average value for each measure in a layer I heterotopion or non-heterotopic layer II/III region from a single animal. Data represent 493 layer I heterotopic neurons and 594 layer II/III neurons from *n* = 7 mice. Wilcoxon matched-pairs, signed rank non-parametric test was used for all three quantifications; ns, no significance; HTP, heterotopia. Descriptive statistics are given in Supplementary File 1. **e**, *In vivo* z-projection of the same GCaMP6f-transfected heterotopion displayed in (**a**), captured at a different depth, showing neurons that were imaged during brief applications of whisker stimulation. **f**, Baseline calcium dynamics (left column) and calcium fluctuations (middle and right columns) of individual heterotopic neurons in response to stimuli (denoted by solid blue line in two trials) obtained from the awake, head-fixed mouse imaged in (**e**). Experimental observations in (**e**, **f**) were replicated in three mice. All data and images from (**a-f**) were obtained from P50-60 mice with heterotopia induced at E15, and virally transfected with AAV-GCaMP6f at P21-P30. Some neuronal cell body and axonal labelling by pCAG-TdTomato (red) is the result of E15 IUE-mediated transfection. Scale bars, 50 µm (**a**), 25 µm (**b**), and 50 µm (**e**).

## DISCUSSION

Here we describe an inducible model that allows the visualization of focal heterotopia at cellular resolution in the live animal. By mechanically puncturing the cortex as part of the IUE procedure we induce a targeted layer I heterotopion that can be repeatedly imaged *in vivo*. Pairing this methodology with intravital fluorescence and label-free (SCoRe) microscopy of neuronal cell bodies, axons and myelin, as well as time lapse axonal and calcium imaging^22,23,25,34,44^, enables detailed studies of heterotopia in the intact brain.

Many animal models of focal heterotopia have been described however most involve subcortical or intrahippocampal heterotopia^13,14,17,19,45^, which are not easily amenable to intravital optical imaging due to their distance from the cortical surface. Moreover, other animal models of more superficially-occurring heterotopia^15,16,18,21,46^ have not been imaged successfully at high resolution *in vivo* mainly as a result of their temporally and spatially unpredictable nature and/or lack of visible cell transfection. Our methodology improves upon these limitations by employing minimally invasive microinjections to both induce layer I heterotopia and deliver plasmid DNA or viral vectors for cellular genetic manipulation in the live mouse.

Using our model, we revealed several striking characteristics of neocortical heterotopia. First, we discovered that axonal projections within layer I heterotopia formed swirled and concentric morphologies specifically around heterotopic neuronal clusters. Interestingly there was no evidence of glial scarring or mechanical tissue barriers surrounding the heterotopia, suggesting that the directional changes in axon pathfinding were instead likely mediated by disruptions in the precise balance of repulsive and attractive gradients of local axon guidance cues^47–51^. These axon pathfinding abnormalities led us to investigate whether the developmental timing of heterotopia induction could modulate their formation. We found that whereas embryonically-induced heterotopia always displayed distinct axon guidance abnormalities, similar defects were not observed in postnatally-induced heterotopia. This surprising finding suggested the presence of a critical gestational time period during which cortical axon pathfinding is uniquely sensitive to local environmental disturbances. The distinct responsiveness of axons during this critical period could be mediated by the enhanced expression of axon growth cone receptors that increase the ability of axons to extend toward guidance cues^52,53^. However, developmentally modulated changes in the signaling gradients of guidance cues themselves could also play a role^54^.

In addition to aberrant axonal organization, we used label-free SCoRe *in vivo* imaging to discover restricted hypermyelination specifically within the heterotopia. This focal hypermyelination was due to an increased production of myelinating oligodendrocytes that deposited myelin sheaths primarily along the aberrantly projecting and concentric axons. Interestingly, the hypermyelination was only observed in embryonically-induced but never in postnatally-induced heterotopia. Similar to the critical period for disrupted axonal patterning, these data suggest that the myelination of embryonically-induced heterotopia is instructed by developmentally regulated molecules or biophysical cues, which could accelerate the local production of myelin by stimulating oligodendrocyte precursor cell recruitment and/or differentiation. Myelin formation is influenced by axon caliber^55^ and genetic manipulations increasing axon caliber can trigger oligodendrocyte precursor cell proliferation and the ensheathment of classically unmyelinated fibers^56^. Thus, axon caliber could serve as one mechanism that differs between the aberrantly projecting axons found in embryonically-induced heterotopia compared to adjacent non-myelinated cortical axons. Other potential causes for the localized hypermyelination of heterotopic axons could include cell surface markers or molecular identity. Specific subpopulations of excitatory^57^ and inhibitory^58^ neurons exhibit variable myelination patterns, seemingly not directly linked to axon caliber, but instead potentially due to their differential expression of adhesion molecules or altered patterns of neuronal activity. Our model provides a novel means to test these possibilities in future studies.

Despite the striking axon and myelin abnormalities associated with the induced heterotopia, we found no evidence of spontaneous epileptiform activity occurring in heterotopic regions or nearby non-heterotopic cortices of awake animals, consistent with previous studies^40,59,60^. Moreover, spontaneous calcium fluctuations and relative synchrony of firing between neurons in layer I heterotopia and neighboring layer II/III cortex were similar. This lack of observable difference in activity may reflect the similar neural composition of both brain areas, which could enable a similar balance of excitatory and inhibitory inputs to these regions. Indeed, we identified Cux1^+^ cells, GABAergic neurons and GABAergic puncta in layer I heterotopia that are also prevalent in layers II/III^37,61–63^. A second, non-mutually exclusive possibility is that more metabolically demanding conditions are required to trigger aberrant activity in mice with induced heterotopia. Consistent with this hypothesis, the application of normally subthreshold doses of convulsant drugs can provoke epileptiform activity in mice with heterotopia *in vitro*^64^ and *in vivo*^59,64,65^. Other animal models of heterotopia, such as the *tish* mutant^13^ and Ihara’s genetically epileptic rat^19^ have documented spontaneous epileptiform activity *in vivo*. Reasons for this difference from our findings remain unclear, however they may be related to disparities in the number, location or size of heterotopia, or unknown off-target effects of the inherited mutations themselves in the genetic models.

There is little known about the functional dynamics of individual heterotopic neurons during behavioral stimulation in the live animal. Using time-lapse calcium imaging, we revealed that heterotopic neurons display robust responses following brief whisker stimulation in awake mice, consistent with the functional connectivity implicated by previous anatomical, electrophysiological, and behavioral studies^13,40,59, 66–69^. This functional connectivity suggests that, even though we did not detect aberrant spontaneous activity within heterotopia, these neurons are integrated into the local neural network and are potentially capable of altering neural network function at baseline or under metabolically demanding conditions.

We describe an inducible model of heterotopia amenable to intravital visualization that closely mimics the neural, glial, axonal, and myelin profiles of those occurring spontaneously in layer I^28,33^. Building upon previous models, this system employs *in utero* microinjections to directly provoke the disruptions in pial basement membrane and radial glial scaffolding thought to culminate in the neural migration defects leading to their development^70^. Importantly, this model allows imaging of the downstream effects of this disruption using multiple optical, functional, and molecular probes for different cell subtypes^22,23,25,34,44^, adding a powerful new tool for the study of cortical malformations in the live mammalian brain.

## MATERIALS AND METHODS

### Animals

All experimental approaches and procedures were conducted in accordance with Yale University Institutional Animal Care and Use Committee regulations. Timed pregnant outbred CD1 mice were purchased from Charles River Laboratories, Inc, with the first 24 hours of postnatal life designated as P0. We included both male and female mice, aged to P30 – P60, for this study. For some birth dating experiments as described in the text, pregnant dams were given a single intraperitoneal (i.p.) injection of EdU (30 µg g^-1^ body weight) at E11.5, prior to IUE surgery at E15. Litters were kept in individual ventilated cages until weaning age (P21), after which mice were housed in single-sex groups with 2 – 5 animals per unit. Cages were maintained in temperature-controlled facilities with 12-hour light/12-hour dark cycles.

### In utero intracranial injection, electroporation and viral infection

Embryonic cortical injections were performed as part of the IUE procedure during needle insertion into the lateral ventricle^23,71^. Embryos were injected once unilaterally between embryonic day 14 – 17 (E14 - E17) for all electroporations, as specified in the text. Intracranial injections were not performed at gestational ages below E14 or over E17 due to the technical challenges associated with maintaining embryo viability. To keep consistent experimental design parameters, however, all quantifications for *in vivo* and fixed tissue analyses used only E15-injected embryos.

All *in utero* injections were targeted toward prospective somatosensory cortices. Injection solutions included the following plasmid and viral components for neuronal labelling: pCAG-tdTomato (based on Addgene plasmid 11150) and rAAV8-hSyn-eGFP (UNC Vector Core, Lot AV5075D; titer 3.9e12 GC ml^-1^). Injection solutions contained either pCAG-tdTomato (1.5 µg µL^-1^ final concentration) only, or both pCAG-tdTomato (1.5 ug µl^-1^ final concentration) and AAV8-hSyn-eGFP (final titer 3.9e11 GC ml^-1^) in saline solution. All injection solutions contained Fast Green FCF dye (TCI; 2 mg ml^-1^) to facilitate their visual tracking during the injection procedure. Procedures for *in utero* electroporation have been previously described^23,71^. Briefly, timed pregnant CD1 dams were anesthetized with a saline solution containing both ketamine (100 - 120 mg kg^-1^) and xylazine (10 - 12 mg kg^-1^), delivered i.p. Following induction of deep surgical anesthesia, midline incisions (1¼ inch) were made into the abdominal skin and muscle wall to access the underlying uterus. Pulled glass capillary needles (10 µL Drummond Scientific Glass Capillaries, Cat# 3-000-210-G; pulled to ∼ 50 µm diameter at tip and ∼ 125 µm diameter at 1mm above tip) were then used to puncture the uterine wall and deliver (Picospritzer II, General Valve) 0.5 µl of injection solution into the lateral ventricle of individual embryos. Each embryo was injected only once and subsequently electroporated using BTX tweezertrodes aimed at the somatosensory cortex of the injected hemisphere. All electroporations were conducted using four 50 ms, 50 V electrical pulses, delivered at 1 second intervals (BTX Harvard Apparatus 8300 pulse generator).

### Postnatal intracranial viral injection

Intracranial injections were performed on P0 pups from timed pregnant CD1 dams that were naïve to the *in utero* electroporation procedure. Injection solutions included one of the following viruses to mark injected regions: rAAV8-hSyn-eGFP (UNC Vector Core, Lot AV5075D; titer 3.9e12 GC ml^-1^) and rAAV2-CaMKIIa-mCherry (UNC Vector Core, Lot AV4377d, 3.8e12 titer GC ml^-1^). P0 intracranial injections were performed essentially as previously described^72^. Briefly, neonates were cryoanesthetized^73^ within 24 hours following birth. After confirming loss of voluntary movement, neonates were placed in the prone position on a polymer cooling block. A pulled glass capillary needle (10 µl Drummond Scientific Glass Capillaries, Cat# 3-000-210-G; pulled to ∼ 75 µm diameter at tip and ∼ 325 µm diameter at 4 mm above tip), advanced 4mm past the dorsal scalp at a 90° angle, was then used to deliver into the brain parenchyma 1 µl of virus solution (diluted 1:10 in PBS and 2 mg ml^-1^ Fast Green FCF, TCI from stock). Each neonate received only one unilateral intracranial injection targeted over somatosensory cortices. Injected neonates were then rewarmed on a heating pad and placed back with their biological mother. All P0-injected mice were sacrificed for analysis at P30, with quantifications and analyses performed in fixed tissue to circumvent the obstructive meningeal scarring and parenchymal adhesions associated with P0-injections on *in vivo* imaging.

### Postnatal subarachnoid viral injection

Injection solutions were prepared using AAV9-Syn-GCaMP6f-WPRE-SV40 (Penn Vector Core, Lot CS1001; titer 7.648e13 GC ml^-1^) virus diluted 1:100 in PBS and Fast Green FCF (TCI; 2 mg ml^-1^). P21-P30 mice that had previously been electroporated *in utero* were anesthetized by intraperitoneal injections of ketamine (100 mg kg^-1^) and xylazine (10 mg kg^-1^). The scalp was shaved, cleaned, and then incised to expose the underlying bone. A high speed drill was next employed to introduce a small burr hole (∼ 0.75 mm diameter) over the transfected hemisphere, taking care to avoid cortical regions suspected to have been directly punctured as part of the *in utero* electroporation procedure. The underlying dura was gently detached, and a 12 µL volume of injection solution (prepared as described above) was infused into the subarachnoid space to achieve viral transfection of both layer I heterotopic and layer II/III neurons via topical cortical application. The scalp incision was then closed with sutures. Cranial windows for *in vivo* imaging were prepared over injected cortical hemispheres 3 – 4 weeks following the subarachnoid AAV infusion.

### Cranial window surgery and in vivo imaging

All *in vivo* imaging was performed using cranial windows^26^. Briefly, mice were anesthetized using ketamine (100 mg kg^-1^) and xylazine (10 mg kg^-1^), delivered via i.p. injection. The dorsal skull surgical field was shaved and cleaned, and a ∼4 mm diameter circular region of skull and dura mater was excised from the injected hemisphere. A #0 transparent glass coverslip was then gently implanted on top of uncovered pial surface to serve as the cranial window. Glue and dental cement were applied to secure the window to surrounding skull bone.

To detect neuronal cell bodies *in vivo* as described in the text, the fluorescent membrane-permeable probe NeuO (NeuroFluor, Stemcell Technologies Cat# 01801, diluted 1:25 in PBS) was applied to exposed cortex for 20 min followed by a 1-2 min PBS rinse, before the placement of the #0 glass coverslip during the cranial window surgery. In some cases, 100 µL Evans blue (TCI; 1 mg ml^-1^) was injected intravenously after the cranial window surgery to label the cortical vasculature.

Except for GCaMP6f calcium imaging experiments, all *in vivo* imaging studies used mice that were anesthetized via i.p. ketamine and xylazine injection. *In vivo* imaging of anesthetized mice was performed at P30 immediately after cranial window surgery. All intravital GCaMP6f calcium imaging was carried out in P50 - P60 awake head-fixed mice starting four hours after arousal from cranial window surgical anesthesia.

Confocal *in vivo* images of previously injected cortical areas with or without the needle tract sites were acquired using a 20X water immersion objective (Leica, 1.0 NA) on a Leica SP5 upright laser scanning microscope. Spectral confocal reflectance (SCoRe) imaging to detect myelinated axon segments was performed as previously described^25,26^ by capturing the simultaneously reflected light signals from 488 nm, 561 nm, and 633 nm multi-wavelength laser excitation outputs. Single-photon laser outputs were tuned to the following excitation wavelengths for fluorescence imaging: 488 nm for GFP and NeuO; 561nm for tdTomato and mCherry; 633nm for Evans Blue. Sequential imaging was employed to minimize overlap between SCoRe reflection and individual fluorescence emission signals for all *in vivo* confocal imaging experiments.

Needle tracts from injections performed *in utero* were identified by the abrupt changes in orientations of labelled dendritic and axonal processes around breaks in the cortical surface *in vivo*. Since needle tracts identified in this manner always displayed ectopic neural clusters of similar expanse in layer I, the outer boundaries of ‘heterotopia’ *in vivo* were defined as the layer I needle tract borders for all quantifications. Ipsilateral, non-injected cortical areas were defined as layer I regions at least 150 µm away from a discernable injection site border. Heterotopia and ipsilateral control regions were imaged using identical laser output and image acquisition configurations that were determined for each experimental data set. Confocal z-stacks were acquired at 1024 × 1024 pixel resolution starting from the pia through depths of up to 120 µm below the cortical pial surface. In some cases, time lapse imaging of *in utero-*induced heterotopia was also performed at 512 × 512 pixel resolution.

For some GCaMP6f calcium imaging experiments that involved visualization of deeper cortical regions in layers II/III as indicated in the text, time-lapse fluorescence images were acquired using a 20X water immersion objective (Zeiss, 1.0 NA) on a Prairie Technologies two-photon microscope fitted with a mode-locked, tunable Spectra Physics Mai-Tai laser. In these experiments, the two-photon laser was tuned to 920nm for excitation of both GCaMP6f and TdTomato fluorophores. All two-photon time-lapse imaging was performed at 512 × 512 pixel resolution.

### In vivo image processing and quantification

Except for GCaMP6f calcium imaging experiments which were conducted using P50 - P60 mice, all intravital imaging studies were performed using P30 mice, with quantifications carried out on those that had previously been injected at E15. All *in vivo* images were processed and quantified using ImageJ/FIJI.

For SCoRe density quantifications, we analyzed single z-sections located 10 µm deep to the cortical pial surface from both heterotopic and ipsilateral control regions. SCoRe density values were assessed using a custom-built, automated thresholding and binarization macro in Image/FIJI, with Robust Automatic Threshold parameters set to noise = 25, lambda = 3, min = 31. Randomly selected, equally sized regions of interest (ROIs) were used to determine SCoRe density values within the centers and edges of the heterotopia versus control regions. A ‘heterotopion center’ was defined as the circular region with a radius extending from the needle tract center to 1/3 of the radius of the needle tract. A ‘heterotopion edge’ was defined as the needle tract concentric circular region just outside the ‘heterotopion center’, extending from the ‘heterotopion center’ outer edge to the outermost border of the needle tract. Average SCoRe densities were determined for the heterotopion center, heterotopion edge and control area for each mouse (*n* = 8 mice). Statistical analyses were carried out using Wilcoxon matched pairs, signed-rank non-parametric tests.

For NeuO dye-labelled cell body density quantifications, data were analyzed from the first 75 µm of cortex deep to the pial surface of both heterotopic and control regions in layer I. Equal-sized volumes (50 µm^3^) were randomly selected from the superficial cortical z-stacks captured from both heterotopic and control regions. The number of NeuO^+^ cell bodies was manually counted in each volume. Average NeuO^+^ cell body densities were then determined for the heterotopic and control region of each mouse (*n* = 6 mice). Statistical analyses were performed using Wilcoxon matched pairs, signed-rank non-parametric tests.

For quantification of neuronal calcium dynamics in heterotopia vs. adjacent non-injected cortical areas, we used the two-photon microscope to capture 150 × 150 µm field of view (FOV) time lapse images from both layer I heterotopia and non-injected ipsilateral layer II/III regions. Time lapse imaging (512 × 512 pixel resolution; 2Hz) was performed in awake, head-fixed mice during a 2-hour time window that started 4 hours after their arousal from cranial window surgical anesthesia. For heterotopic regions, FOVs were randomly selected from within the first 75 µm below the cortical surface in layer I. For layer II/III regions, FOVs were randomly selected from between 120 to 175 µm below the cortical surface. Six to eight separate FOVs (exactly half from layer I heterotopia and half from ipsilateral layer II/III regions) were imaged in each mouse, with each FOV recorded for 120s per trial for three trials. The order in which FOVs were acquired from heterotopia and layer II/III regions was alternated between mice. Time series analyses were performed using ImageJ/FIJI. Prior to quantifications, TurboReg plugin in ImageJ/FIJI was used to align all time series images in the XY plane.

To quantify GCaMP6f fluorescence changes in individual cells, ROIs were manually selected to encapsulate neuronal cell bodies. GCaMP6f^+^ cell bodies that exhibited tdTomato labelling from the IUE procedure were excluded from all analyses. Approximately 10 - 25 GCaMP6f^+^ cells were analyzed in each FOV. Baseline GCaMP6f fluorescence intensity values (F) for each cell were defined in each trial as the average of the lowest 20% of the recorded values per trial, with fluctuations from this baseline denoted as ΔF/F. A spike event was defined in each trial as a ΔF/F > 0.5. Spike event frequencies were averaged across 3 trials for each cell in both layer I heterotopia and layer II/III regions for *n* = 7 mice. Statistical analyses were performed using Wilcoxon matched pairs, signed-rank non-parametric tests.

For quantifications of the global synchronization index reflecting the relative coordination of GCaMP6f activity, analyses were performed using Fluorescence Single Neuron and Network Analysis Package (FluoroSNNAP)^74^. Briefly, this semi-automated software implements a correlation matrix-based algorithm^74–76^ to compute the normalized global synchronization indices (ranging in value from 0 to 1) for time series calcium imaging data of neural populations. The highest value, 1, signifies entirely synchronized firing throughout an identified cluster of neurons, whereas 0 indicates the total absence of synchrony. Synchronization cluster analyses were performed using template-based calcium event detection parameters set to detection threshold = 0.85, minimum size of synchronization clusters = 2, and surrogate resampling = 20. A cluster of neurons was defined as the GCaMP6f^+^ cell population within one FOV. Global synchronization indices were averaged across three trials for each FOV in both layer I heterotopia and layer II/III regions for *n* = 7 mice. Statistical analyses were performed using Wilcoxon matched pairs, signed-rank non-parametric tests.

### Tissue processing and immunohistochemistry

At P30, mice were deeply anesthetized using ketamine and xylazine, and then perfused transcardially with 4% paraformaldehyde in phosphate buffered saline (PBS). Harvested brains were post-fixed overnight using the same solution at 4 °C, and then vibratome sectioned (coronal; 75 µm thickness) for fixed tissue analysis.

To identify brain regions that had been injected as part of the IUE procedure, a fluorescence microscope was next used to examine the sections for visibly disrupted layer I regions showing transfected neurons and/or neuronal processes, similar to what has been previously described^46^. The disrupted regions often revealed cellular clusters containing eGFP^+^ neurons that protruded past the pial surface, which were never observed in intact cortical regions or in non-injected mice. Sections containing visibly disrupted layer I cortical cytoarchitecture indicative of injection-associated trauma were then selected for further processing. Immunohistochemistry, performed as described below, was used to confirm the presence of layer I heterotopia at all identified IUE injection sites. Coronal sections from P0-injected mice were screened in a similar manner for injected areas, which were identified by virally transfected axonal fibers and neuronal cell bodies concentrated around visible needle tracts (data not shown).

Prior to immunostaining, all free-floating sections were heated to 95 °C for 30 minutes in 50 mM sodium citrate buffer (0.05% Tween-20, pH 6.0) for antigen retrieval and elimination of endogenous eGFP expression. After rinsing the sections in PBS at room temperature, EdU labelling was next carried out in some experiments as specified by the Click-iT EdU Alexa Fluor-647 Imaging Kit protocol (Cat# C10340). Before proceeding to antibody staining, all tissues were pre-incubated for 1-2 hr in 0.1% Triton X-100 and 5% Normal Goat Serum (NGS; Jackson Immunoresearch, Cat# 005-000-121) in PBS at room temperature. Slices were then incubated with primary antibody for 1.5 hrs - 2 days as needed in 0.1% Triton X-100 and 5% NGS in PBS at 4 °C. The primary antibodies used were: rabbit anti-NeuN (Abcam, Cat# ab177487, 1:3000), mouse anti-NeuN (Abcam, Cat# ab104224, 1:1000), mouse anti-Myelin CNPase (clone SMI 91, Biolegend, Cat# 836404, 1:1000), rabbit anti-Cux1 (Novus Biologicals, Cat# NBP2-13883, 1:100), mouse anti-Tle4 (E-10) Alexa Fluor 647 (Santa Cruz Biotechnology, Cat# sc-365406 AF647, 1:100), mouse anti-GAD-67 (EMD Millipore, Cat# MAB5406, 1:1000), rabbit anti-Iba1 (Wako, Cat# 019-19741, 1:600), mouse anti-Aldh1l1 (clone N103/39, NeuroMab, Cat# 75-140, 1:500), chicken anti-Neurofilament NF-H (EnCor, Cat# CPCA-NF-H, 1:500), chicken anti-GFP (Abcam, Cat# 13970, 1:500), and rabbit anti-Myelin Basic Protein (MBP; Abcam, Cat# 40390, 1:1000). Following primary antibody incubation, slices were washed in PBS and then incubated with Alexa Fluor dye-conjugated secondary antibodies of the appropriate host species at 1:600 dilution in 0.1% Triton X-100 and 5% NGS in PBS for 1 - 2 days at 4 °C. After secondary antibody incubation, sections were washed again in PBS and incubated with 2.5 µg mL^-1^ 4′,6-diamidino-2-phenylindole (DAPI) in PBS for 15 min at room temperature to counterstain cell nuclei. Following an additional subsequent wash, stained sections were mounted with 25% mounting media solution (Dako Ultramount, Cat# S1964; diluted 1:4 in PBS) onto glass slides for imaging.

### Fixed tissue imaging and quantification

Images of stained sections were collected using a Leica SP5 upright confocal laser scanning microscope. Lower magnification views of analyzed sections for presentation were acquired using 10X and 20X Leica objectives. All fixed tissue images for quantification were captured through a 40X Leica water immersion objective. For immunohistochemical analyses of *in utero*-injected mice, ‘heterotopia’ were defined as the layer I cortical regions demarcated by ectopic NeuN^+^ cell body clusters identified at IUE needle tract sites. For P0-injected mice, because the heterotopia associated with needle tracts were more variable in size, ranging from several cells to 350 µm in diameter (data not shown), data for these injected areas were obtained from 300 µm X 90 µm FOVs centered on visible needle tracts that were within the first 100 µm below the pial surface. Control, non-injected regions were specified on corresponding contralateral cortices at similar mediolateral distances from the midline. Identical laser output and image acquisition configurations were used to capture images of both heterotopia and contralateral control regions throughout each immunostaining set.

All data for Aldh1l1, Iba1, CNPase, and GAD-67 cell density quantifications were acquired from the most superficial 100 µm of cortical tissue of both heterotopic and contralateral homologous control regions in layer I. The number of cell bodies confirmed by DAPI labelling that were positive for each marker was manually counted in randomly selected volumes within z-stacks acquired from both regions. The volumes were kept at identical dimensions between heterotopia and contralateral control regions for each experiment. For GAD-67 immunostaining, the presence of neuronal cell bodies was also confirmed by NeuN labelling. Two coronal sections containing heterotopia were analyzed per animal, with each immunostaining set comprising *n* = 6 animals. Statistical analyses were performed using Wilcoxon matched pairs, signed-rank non-parametric tests.

Data for CNPase density quantifications were acquired from the most superficial 100 µm of cortical tissue in both heterotopia and contralateral control regions. CNPase intensity was assessed using an automated thresholding and binarization plugin in Image/FIJI (Robust Automatic Threshold parameters set to noise = 1, lambda = 2, min = 208) for equally-sized ROIs that were randomly selected from within heterotopic and contralateral control z-projections (15 µm thickness). CNPase density values for each section were determined by subtracting the automated measurements made in additional background regions from those made in assessed heterotopic and control regions. Two coronal sections containing heterotopia were analyzed per animal (*n* = 6 animals). Statistical analyses were performed using Wilcoxon matched pairs, signed-rank non-parametric tests.

To examine the neuronal composition of the embryonically-induced heterotopia, coronal sections were stained for NeuN, DAPI and one of the other following stains: Cux1, Tle4, or EdU as indicated in the text. Data were acquired from within the most superficial 100 µm of the neocortex for layer I heterotopia, and across the entire cortical wall for contralateral homologous control regions. For quantification of layer I heterotopia, equally sized VOIs were randomly selected from within the first 100 µm below the pial surface. For quantification of control regions on the contralateral hemisphere, the cortical wall was partitioned into 11 equal volume bins extending from the pial surface to white matter, with Bin1 encompassing layer 1 and Bin 11 adjoining white matter at its lower boundary. For each heterotopic VOI and control bin, the 1) number of NeuN^+^ cells and 2) proportion of NeuN^+^ cells that were also Cux1, Tle4, or EdU positive was manually quantified in ImageJ. Cells were identified as EdU positive if > 50% of their nuclear volume, defined by DAPI, was occupied by EdU. Two sections were analyzed per mouse, with sample sizes for each group denoted in the text and figure legends.

The sites of injection analyzed in this study were positioned in somatosensory cortices and immediately adjacent areas in both male and female mice. Given the minimal variation in the morphological appearances of needle tracts between male and female mice and across different cortical areas, we pooled together these variables in our analyses. No animal subjects or experimental data points were excluded from analysis, and the nature of our study did not require subject randomization or experimenter blinding. No statistical methods were used to predetermine sample sizes, although our sample sizes are comparable to those published and generally accepted in the field. GraphPad Prism 7 was utilized for all statistical analyses, and Wilcoxon matched pairs signed-rank non-parametric tests were used to determine statistical significance (declared for p-values below 0.05; two-tailed) because a normal distribution of differences between the paired data could not be assumed. Data are reported as mean ± s.e.m., unless otherwise noted.

## ACKNOWLEDGEMENTS

We thank F. Chen for his guidance *in utero* electroporation techniques. This study was funded by grants from the U.S. National Institutes of Health (R01NS089734, R21NS088411 and R21NS087511 to J.G.; R00NS099469 and P20GM113132 to R.A.H.; T32NS007224 and T32GM007205 to A.M.L.), Donors Cure Foundation (New Vision Award to R.A.H.), and National Multiple Sclerosis Society (#RR-1602-07686 to J.G.).

## AUTHOR CONTRIBUTIONS

A.M.L. and J.G. conceived the initial project. A.M.L., R.A.H, and J.G. designed the experiments. A.M.L. performed all experiments, analyzed the data, and prepared the figures. A.M.L., R.A.H. and J.G. wrote the manuscript.

---

**Figure 3–figure supplement 1.**
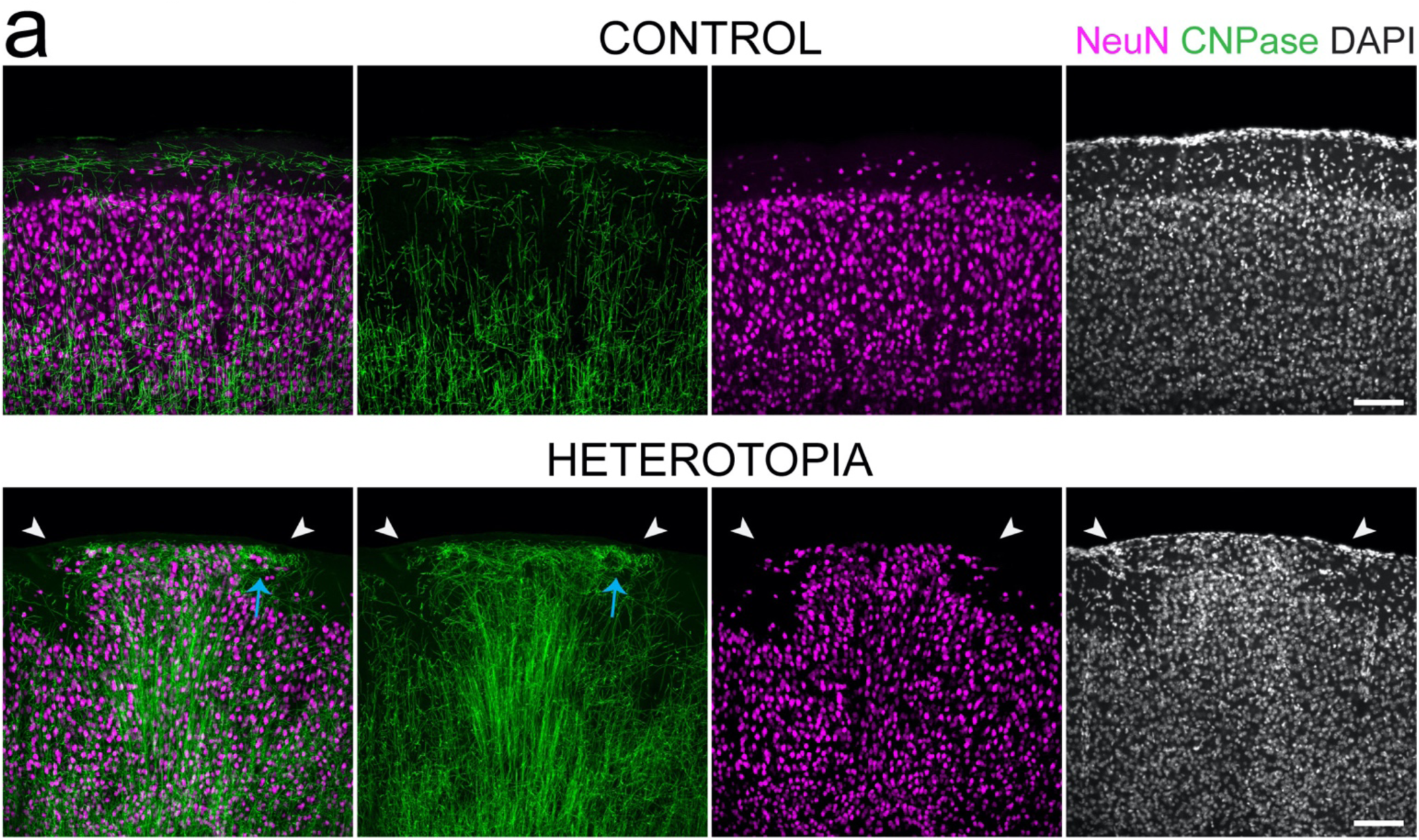
Myelinated axons project aberrantly around heterotopic cells. **a**, High resolution immunostaining showing the winding trajectories followed by CNPase^+^ fibers (bottom, blue arrow) within a layer I heterotopion (NeuN; white arrowheads) in a P30 mouse coronal section. The aberrantly-projecting myelinated fiber bundles are absent from immediately adjacent non-heterotopic cortex and corresponding contralateral control regions (top row). Images are representative of experiments performed in at least six animals. Scale bars, 100 µm.

**Figure 2–figure supplement 1.**
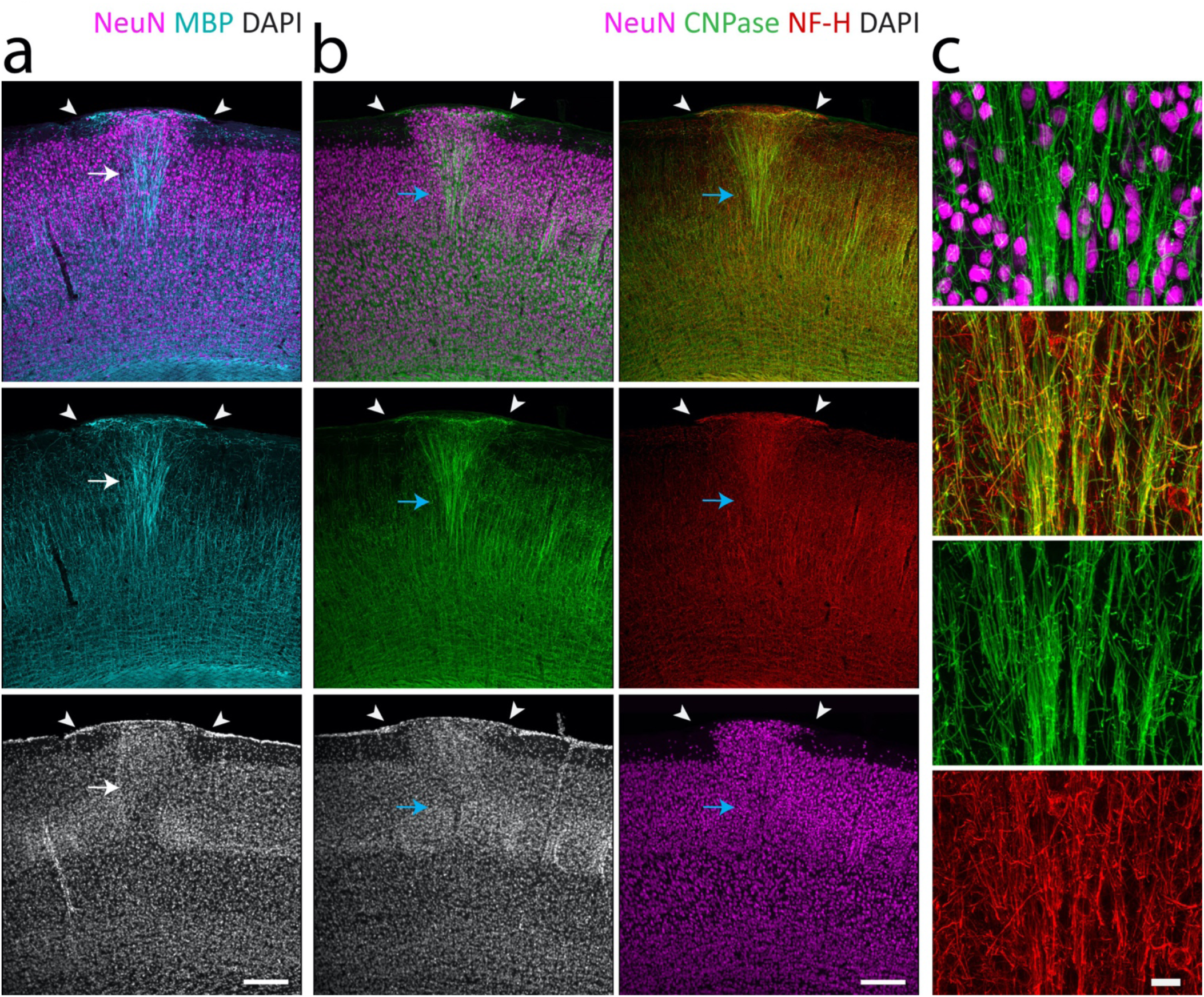
Disrupted myelin patterning and axonal pathfinding occurs in deeper cortical layers. **a,** Myelinated fibers (MBP, cyan) accumulate in thick radially oriented cords (white arrow) beneath a layer I heterotopion (NeuN; arrowheads) in an immunostained P30 mouse forebrain coronal section. **b-c,** Confocal images of a different coronal section from the same heterotopion in (**a**), revealing similar aberrant myelin and axon accumulations as identified by oligodendrocyte CNPase (green) and axonal NF-H (red) immunostaining, respectively. A higher magnification view of the densely packed myelinated axons fasciculating underneath the heterotopion (**b**, blue arrow) is shown in (**c**). Images in (**b**) and (**c**) are of same section shown in Figure 3e. Images are representative of fixed tissue experiments performed in at least three mice. Scale bars, 200 µm (**a**, **b**), and 20 µm (**c**).

**Figure 7–figure supplement 1.**
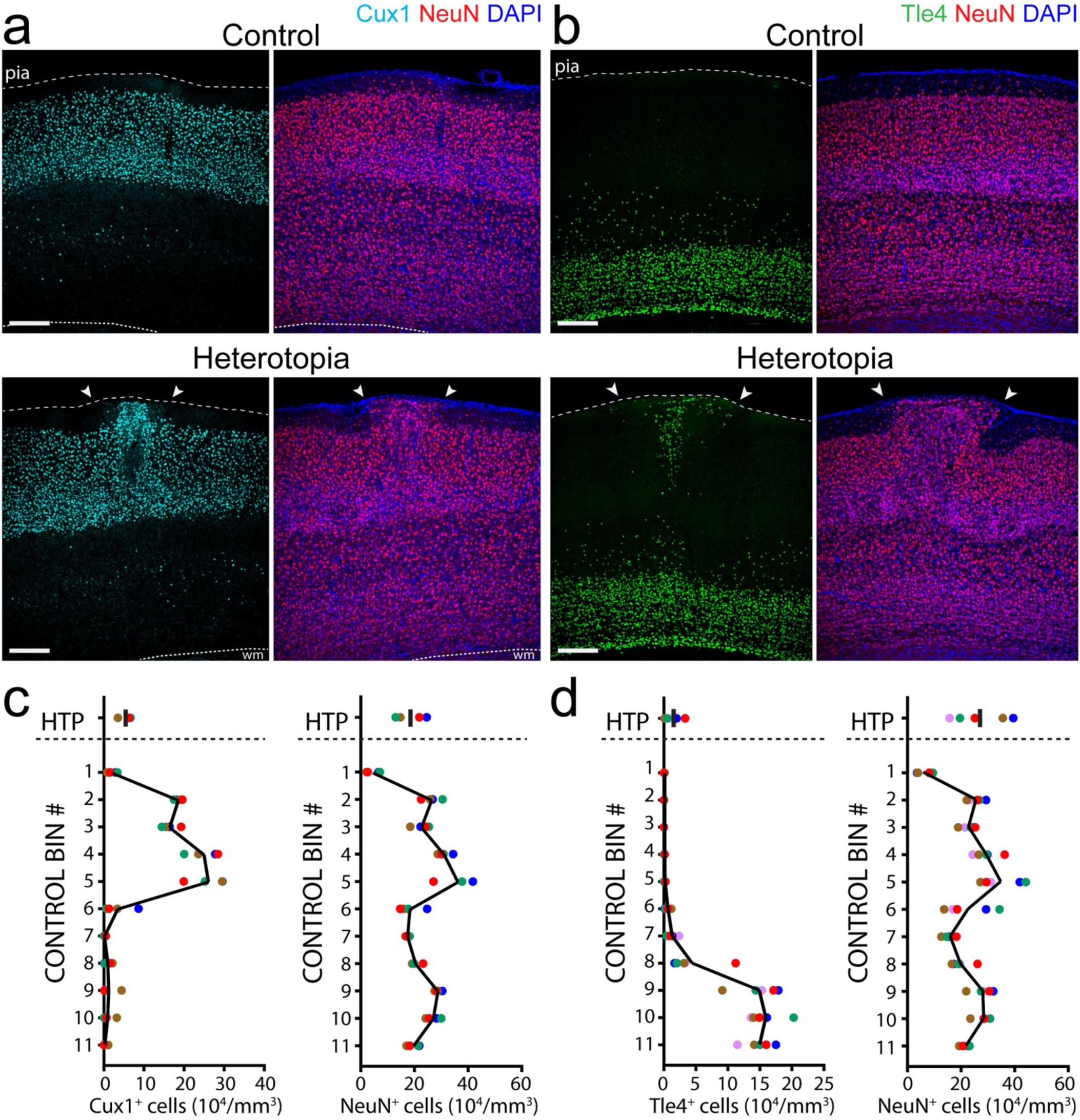
Heterotopic neurons express both deep and superficial cortical layer markers. **a-b**, Additional fixed tissue images captured from the heterotopia displayed in Figures 7b and c, confirming the presence of Cux1^+^ (**a**, cyan) and Tle4^+^ (**b**, green) neuronal cell bodies within the representative heterotopia (bottom images, arrowheads). In corresponding contralateral control regions (top images), Cux1 (left) largely concentrates in superficial cortical layers whereas Tle4 (right) tends to localize toward deeper cortical layers. **c-d**, Quantifications showing the densities of Cux1^+^ (**c**, *n* = 4 animals), Tle4^+^ (**d**, *n* = 5 animals), and NeuN^+^ (**c** and **d**, right) cells in heterotopia and across corresponding contralateral control regions of P30 mice. Each dot corresponds to the layer I heterotopion or control bin of a single animal, with all dots of the same color belonging to the same animal. The black line denotes the mean. Images are representative of experiments performed in at least four animals. WM, white matter; HTP, heterotopia. Scale bars, 200 µm.

**Figure 7–figure supplement 2.**
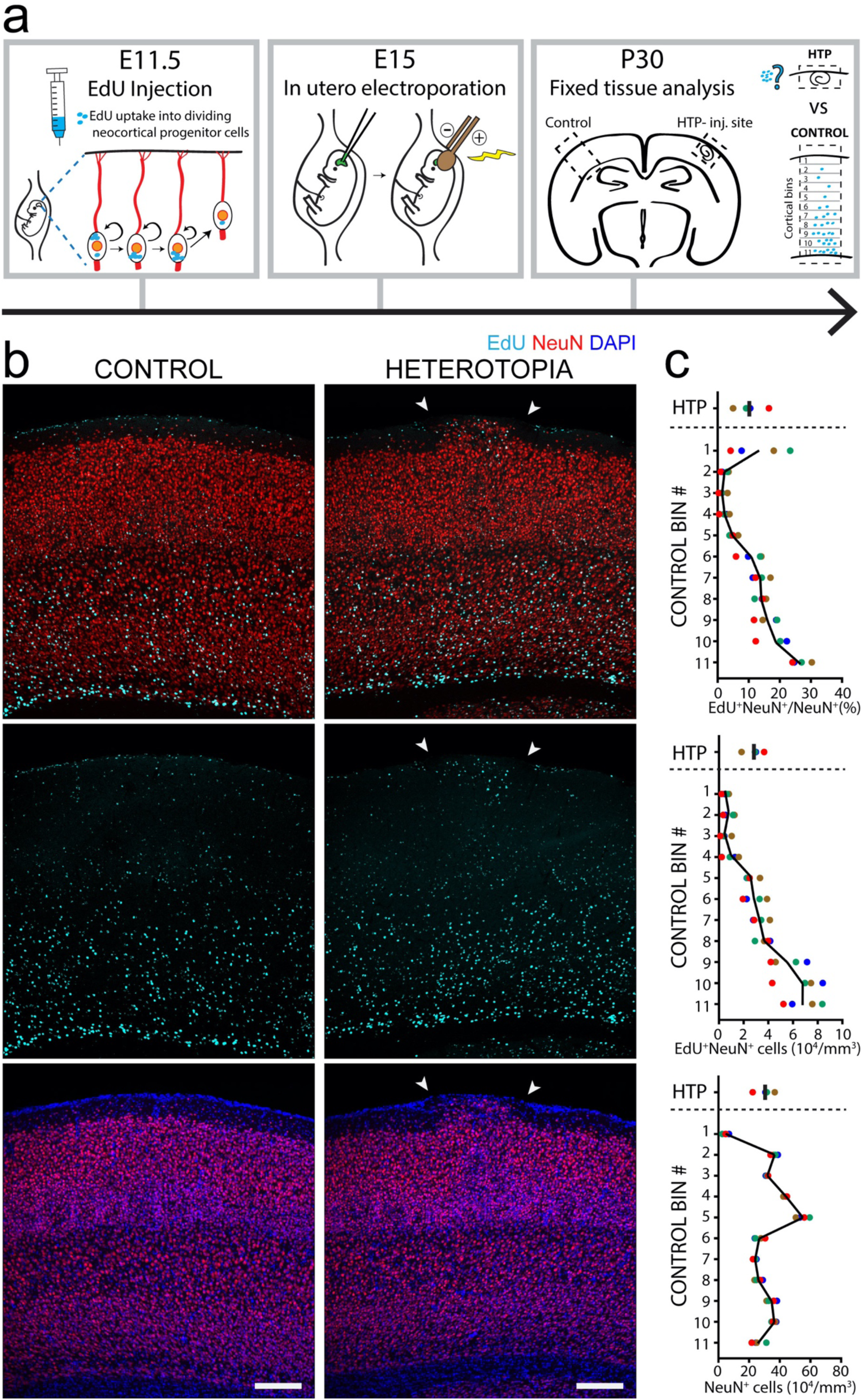
Birth-dating of heterotopic neurons. **a**, Diagram of the pulse labelling strategy for birthdating heterotopic cortical neurons. Dividing cells are labelled by a single EdU injection at E11.5, with heterotopia induced at E15. Mice are sacrificed at P30 for analysis. **b**, Low magnification images showing the EdU^+^ neurons incorporated in a layer I heterotopion (right, arrowheads) of a P30 mouse. In corresponding contralateral control cortex (left column), EdU expression is predominantly concentrated in deeper cortical layers. **c**, Quantifications showing the percentages of NeuN^+^ cells that also express EdU (top), EdU^+^NeuN^+^ cell body densities (middle), and NeuN^+^ cell body densities (bottom) in layer I heterotopia and across corresponding contralateral control cortices of P30 mice. Each dot corresponds to the layer I heterotopion or control bin of a single animal, with all dots of the same color belonging to the same animal (*n* = 4 animals total). Images are representative of the observations made in each mouse brain. The black line denotes the mean. WM, white matter; HTP, heterotopia. Scale bars, 200 µm.

**Video 1. *In vivo* imaging of a layer I heterotopion and adjacent control area.** Confocal z-stacks of a heterotopion (right) and ipsilateral non-injected layer I control region (left) captured from a living P30 mouse using label-free SCoRe (magenta) and fluorescence microscopy, showing the aberrantly projecting myelinated axons and ectopic neurons located inside a layer I heterotopion induced at E15. Neuronal cell bodies and cerebral blood vessels are visualized using fluorescent neuronal dye NeuO (green) and Evans Blue (white), respectively. The density of the horizontally crisscrossing, aberrantly projecting fibers diminishes with increasing cortical depth (indicated at bottom left).

**Video 2. Layer I heterotopia and adjacent layer II/III regions display highly variable spontaneous neuronal calcium transients.** Representative *in vivo* two-photon time-lapse recordings of GCaMP6f-labelled neurons (green) within a layer I heterotopion (right) and an adjacent layer II/III cortical region (left) in an awake, head-fixed P55 mouse. Diverse patterns of spontaneous calcium transients are observed in neuronal cell bodies throughout both imaged regions. Note the absence of obvious epileptiform activity. Images were acquired at 2 Hz from an E15-induced layer I heterotopion. Some neuronal cell body and axonal labelling by pCAG-TdTomato (red) is the result of E15 IUE-mediated transfection.

**Video 3. Heterotopic neurons respond to sensory stimuli.** Two *in vivo* confocal time-lapse recording examples of GCaMP6f-labelled neurons within a layer I heterotopion before, during, and after the application of a brief whisker stimulus, administered one minute into each imaging session. The timing of the stimulus and accompanying movements are indicated by the white circle (positioned at the top left of the embedded video). Neuronal calcium changes are observed upon the application of each stimulus, indicating the responsiveness of the heterotopic neurons to sensorimotor input. Both recordings were obtained from P55 awake, head-fixed mice with layer I heterotopia that were induced at E15 and virally transfected using AAV-GCaMP6f at P21-P30.

**Supplementary Table 1.** Details for all statistical analyses, including sample sizes and p-values.

| Supplementary Table 1 |  |  |  |  |  |
| --- | --- | --- | --- | --- | --- |
| Figure Number | Statistical Test | Sample Size | Sample Definition | P Value | Degrees of Freedom and F/T/z/R/ET C value |
| 1b | Wilcoxon non-parametric matched-pairs signed rank test | 6 | number of mice (paired samples) | p = 0.0313 for NeuO, HTP vs. Control | n/a |
| 2d | Wilcoxon non-parametric matched-pairs signed rank test | 8 | number of mice (paired samples) | SCoRe density, p=0.0078 for HTP edge vs. Control; p=0.0078 for HTP center vs. Control | n/a |
|  | Wilcoxon non-parametric matched-pairs signed rank test | 6 | number of mice (paired samples) | p = 0.0313 for CNPase density, HTP vs. Control. | n/a |
|  | Wilcoxon non-parametric matched-pairs signed rank test | 6 | number of mice (paired samples) | p = 0.0313 for CNPase<sup>+</sup> cell bodies, HTP vs. Control. | n/a |
| 5c | Wilcoxon non-parametric matched-pairs signed rank test | 6 | number of mice (paired samples) | p > 0.9999 for CNPase<sup>+</sup> cell bodies, HTP vs. Control. | n/a |
|  | Wilcoxon non-parametric matched-pairs signed rank test | 6 | number of mice (paired samples) | p = 0.0625 for CNPase density, HTP vs. Control. | n/a |
| 6c | Wilcoxon non-parametric matched-pairs signed rank test | 6 | number of mice (paired samples) | p = 0.8438 for Aldh1L1<sup>+</sup> cell density, HTP vs. Control. | n/a |
| 6f | Wilcoxon non-parametric matched-pairs signed rank test | 6 | number of mice (paired samples) | p = 0.9375 for Iba<sup>+</sup> cell density, HTP vs. Control | n/a |
| 7f | Wilcoxon non-parametric matched-pairs signed rank test | 6 | number of mice (paired samples) | p = 0.5313 for GAD67<sup>+</sup> cell density, HTP vs Control | n/a |
| 8d | Wilcoxon non-parametric matched-pairs signed rank test | 7 | number of mice (paired samples) | p = 0.6875 for spike event frequency, HTP vs Layers II/III | n/a |
|  | Wilcoxon non-parametric matched-pairs signed rank test | 7 | number of mice (paired samples) | p = 0.9375 for variance (s.d.), HTP vs. Layers II/III | n/a |
|  | Wilcoxon non-parametric matched-pairs signed rank test | 7 | number of mice (paired samples) | p = 0.2969 for global synchronization index, HTP vs. Layers II/III | n/a |

## REFERENCES

1. Kaufmann WE, Galaburda AM. Cerebrocortical microdysgenesis in neurologically normal subjects: a histopathologic study. Neurology. 1989;39(2 Pt 1):238–244. doi:10.1212/WNL.39.2.238

2. Kasper BS, Stefan H, Buchfelder M, Paulus W. Temporal Lobe Microdysgenesis in Epilepsy Versus Control Brains. J Neuropathol Exp Neurol. 1999;58(1):22–28. doi:10.1097/00005072-199901000-00003

3. Meencke HJ, Veith G. Migration disturbances in epilepsy. Epilepsy Res Suppl. 1992;9:31–39; discussion 39-40. http://www.ncbi.nlm.nih.gov/pubmed/1285910.

4. Schulze KD, Braak H. Hirnwarzen. Zeitschrift fur Mikroskopisch-Anatomische Forsch - Abteilung 2. 1978;92(4):609–623.

5. Sisodiya SM. Malformations of Cortical Development: Burdens and Insights from Important Causes of Human Epilepsy. Vol 3. Lancet Publishing Group; 2004:29–38. doi:10.1016/S1474-4422(03)00620-3

6. Barkovich AJ, Gressens P, Evrard P. Formation, maturation, and disorders of brain neocortex. Am J Neuroradiol. 1992;13(2):423–446. http://www.ncbi.nlm.nih.gov/pubmed/1566709.

7. Galaburda AM, Kemper TL. Cytoarchitectonic abnormalities in developmental dyslexia: a case study. Ann Neurol. 1979;6(2):94–100. doi:10.1002/ana.410060203

8. Hardiman O, Burke T, Phillips J, et al. Microdysgenesis in resected temporal neocortex: incidence and clinical significance in focal epilepsy. Neurology. 1988;38(7):1041–1047. doi:10.1212/wnl.38.7.1041

9. Palmini A, Andermann F, Olivier A, Tampieri D, Robitaille Y. Focal neuronal migration disorders and intractable partial epilepsy: Results of surgical treatment. Ann Neurol. 1991;30(6):750–757. doi:10.1002/ana.410300603

10. Kakita A, Hayashi S, Moro F, et al. Bilateral periventricular nodular heterotopia due to filamin 1 gene mutation: Widespread glomeruloid microvascular anomaly and dysplastic cytoarchitecture in the cerebral cortex. Acta Neuropathol. 2002. doi:10.1007/s00401-002-0594-9

11. Farrell MA, DeRosa MJ, Curran JG, et al. Neuropathologic findings in cortical resections (including hemispherectomies) performed for the treatment of intractable childhood epilepsy. Acta Neuropathol. 1992;83(3):246–259. http://www.ncbi.nlm.nih.gov/pubmed/1557956.

12. Nordborg C, Eriksson S, Rydenhag B, Uvebrant P, Malmgren K. Microdysgenesis in surgical specimens from patients with epilepsy: occurrence and clinical correlations. J Neurol Neurosurg Psychiatry. 1999;67(4):521–524. http://www.ncbi.nlm.nih.gov/pubmed/10486403.

13. Lee KS, Schottler F, Collins JL, et al. A genetic animal model of human neocortical heterotopia associated with seizures. J Neurosci. 1997;17(16):6236–6242. doi:10.1523/jneurosci.17-16-06236.1997

14. Bai J, Ramos RL, Ackman JB, Thomas AM, Lee R V., LoTurco JJ. RNAi reveals doublecortin is required for radial migration in rat neocortex. Nat Neurosci. 2003;6(12):1277–1283. doi:10.1038/nn1153

15. Rosen GD, Sherman GF, Richman JM, Stone L V., Galaburda AM. Induction of molecular layer ectopias by puncture wounds in newborn rats and mice. Dev Brain Res. 1992;67(2):285–291. doi:10.1016/0165-3806(92)90229-P

16. Brunstrom JE, Gray-Swain MRR, Osborne PA, Pearlman AL. Neuronal Heterotopias in the Developing Cerebral Cortex Produced by Neurotrophin-4. Neuron. 1997;18(3):505–517. doi:10.1016/S0896-6273(00)81250-7

17. Riggs HE, McGrath JJ, Schwarz HP. Malformation of the adult brain (albino rat) resulting from prenatal irradiation. J Neuropathol Exp Neurol. 1956;15(4):432–447. doi:10.1097/00005072-195610000-00006

18. Sherman GF, Galaburda AM, Behan PO, Rosen GD. Neuroanatomical anomalies in autoimmune mice. Acta Neuropathol. 1987;74(3):239–242. doi:10.1007/BF00688187

19. Amano S, Ihara N, Uemura S, et al. Development of a Novel Rat Mutant with Spontaneous Limbic-Like Seizures. Am J Pathol. 1996;149(1):329–336.

20. Dvořàk K, Feit J, Juránková Z. Experimentally induced focal microgyria and status verrucosus deformis in rats - Pathogenesis and interrelation histological and autoradiographical study. Acta Neuropathol. 1978;44(2):121–129. doi:10.1007/BF00691477

21. Ferrer I, Alcántara S, Catalá I, Zújar MJ. Experimentally induced laminar necrosis, status verrucosus, focal cortical dysplasia reminiscent of microgyria, and porencephaly in the rat. Exp brain Res. 1993;94(2):261–269. http://www.ncbi.nlm.nih.gov/pubmed/8359242. Accessed July 14, 2017.

22. Lo Turco J, Manent JB, Sidiqi F. New and improved tools for in utero electroporation studies of developing cerebral cortex. Cereb Cortex. 2009;19(Suppl. 1):i120–5. doi:10.1093/cercor/bhp033

23. Saito T, Nakatsuji N. Efficient Gene Transfer into the Embryonic Mouse Brain Using in Vivo Electroporation. Dev Biol. 2001;240(1):237–246. doi:10.1006/dbio.2001.0439

24. Tabata H, Nakajima K. Efficient in utero gene transfer system to the developing mouse brain using electroporation: Visualization of neuronal migration in the developing cortex. Neuroscience. 2001. doi:10.1016/S0306-4522(01)00016-1

25. Schain AJ, Hill RA, Grutzendler J. Label-free in vivo imaging of myelinated axons in health and disease with spectral confocal reflectance microscopy. Nat Med. 2014;20(4):443–449. doi:10.1038/nm.3495

26. Hill RA, Li AM, Grutzendler J. Lifelong cortical myelin plasticity and age-related degeneration in the live mammalian brain. Nat Neurosci. 2018;21(May):1–13. doi:10.1038/s41593-018-0120-6

27. Hill RA, Grutzendler J. In vivo imaging of oligodendrocytes with sulforhodamine 101. Nat Methods. 2014;11(11):1081–1082. doi:10.1038/nmeth.3140

28. Jacob H. Die feinere Oberflächengestaltung der Hirnwindungen, die Hirnwarzenbildung und die Mikropolygyrie. Zeitschrift für die gesamte Neurol und Psychiatr. 1940;170:68–84. doi:10.1007/bf02869355

29. Morel F, Wildi E. Dysgénésie nodulaire disséminée de l’écorce frontale. Rev Neurol (Paris). 1952;87(3):251–270.

30. Sherman GF, Stone JS, Press DM, Rosen GD, Galaburda AM. Abnormal architecture and connections disclosed by neurofilament staining in the cerebral cortex of autoimmune mice. Brain Res. 1990;529(1-2):202–207. doi:10.1016/0006-8993(90)90828-Y

31. Ramos RL, Smith PT, DeCola C, Tam D, Corzo O, Brumberg JC. Cytoarchitecture and Transcriptional Profiles of Neocortical Malformations in Inbred Mice. Cereb Cortex. 2008;18(11):2614–2628. doi:10.1093/cercor/bhn019

32. Sherman GF, Galaburda AM, Geschwind N. Cortical anomalies in brains of New Zealand mice: a neuropathologic model of dyslexia? Proc Natl Acad Sci U S A. 1985;82(23):8072–8074. doi:10.1073/pnas.82.23.8072

33. Ramos RL, Siu NY, Brunken WJ, et al. Cellular and axonal constituents of neocortical molecular layer heterotopia. Dev Neurosci. 2014;36(6):477–489. doi:10.1159/000365100

34. Er JC, Leong C, Teoh CL, et al. NeuO: a Fluorescent Chemical Probe for Live Neuron Labeling. Angew Chemie Int Ed. 2015;54(8):2442–2446. doi:10.1002/anie.201408614

35. Humphreys P, Kaufmann WE, Galaburda AM. Developmental dyslexia in women: neuropathological findings in three patients. Ann Neurol. 1990;28(6):727–738. doi:10.1002/ana.410280602

36. Galaburda AM, Sherman GF, Rosen GD, Aboitiz F, Geschwind N. Developmental dyslexia: Four consecutive patients with cortical anomalies. Ann Neurol. 1985;18(2):222–233. doi:10.1002/ana.410180210

37. Molyneaux BJ, Arlotta P, Fame RM, MacDonald JL, MacQuarrie KL, Macklis JD. Novel subtype-specific genes identify distinct subpopulations of callosal projection neurons. J Neurosci. 2009;29(39):12343–12354. doi:10.1523/jneurosci.6108-08.2009

38. Molyneaux BJ, Goff LA, Brettler AC, et al. DeCoN: Genome-wide Analysis of In Vivo Transcriptional Dynamics during Pyramidal Neuron Fate Selection in Neocortex. Neuron. 2015;85(2):275–288. doi:https://doi.org/10.1016/j.neuron.2014.12.024

39. Sorensen SA, Bernard A, Menon V, et al. Correlated Gene Expression and Target Specificity Demonstrate Excitatory Projection Neuron Diversity. Cereb Cortex. 2015;25(2):433–449. doi:10.1093/cercor/bht243

40. Gabel LA, LoTurco JJ. Electrophysiological and morphological characterization of neurons within neocortical ectopias. J Neurophysiol. 2001;85(2):495–505. doi:10.1152/jn.2001.85.2.495

41. Gabel LA. Layer I neocortical ectopia: Cellular organization and local cortical circuitry. Brain Res. 2011;1381:148–158. doi:10.1016/J.BRAINRES.2011.01.040

42. Angevine JB, Sidman RL. Autoradiographic Study of Cell Migration during Histogenesis of Cerebral Cortex in the Mouse. Nature. 1961;192(4804):766–768. doi:10.1038/192766b0

43. Raedler E, Raedler A. Autoradiographic study of early neurogenesis in rat neocortex. Anat Embryol (Berl). 1978;154(3):267–284. doi:10.1007/bf00345657

44. Hill RA, Damisah EC, Chen F, Kwan AC, Grutzendler J. Targeted two-photon chemical apoptotic ablation of defined cell types in vivo. Nat Commun. 2017;8(May):1–15. doi:10.1038/ncomms15837

45. Singh SC. Ectopic neurones in the hippocampus of the postnatal rat exposed to methylazoxymethanol during foetal development. Acta Neuropathol. 1977;40(2):111–116. doi:10.1007/bf00688698

46. Rosen GD, Bai J, Wang Y, et al. Disruption of Neuronal Migration by RNAi of Dyx1c1 Results in Neocortical and Hippocampal Malformations. Cereb Cortex. 2007;17(11):2562–2572. doi:10.1093/cercor/bhl162

47. Bagnard D, Lohrum M, Uziel D, Püschel AW, Bolz J. Semaphorins act as attractive and repulsive guidance signals during the development of cortical projections. Development. 1998;125(24):5043–5053. http://www.ncbi.nlm.nih.gov/pubmed/9811588.

48. Richards LJ, Koester SE, Tuttle R, O’Leary DD. Directed growth of early cortical axons is influenced by a chemoattractant released from an intermediate target. J Neurosci. 1997;17(7):2445–2458. http://www.ncbi.nlm.nih.gov/pubmed/9065505.

49. Métin C, Deléglise D, Serafini T, Kennedy TE, Tessier-Lavigne M. A role for netrin-1 in the guidance of cortical efferents. Development. 1997;124(24):5063–5074. https://www.scopus.com/inward/record.uri?eid=2-s2.0-0031441084&partnerID=40&md5=16c8fc81c15621923b5da3fd9e86e059. Accessed September 23, 2019.

50. Polleux F, Giger RJ, Ginty DD, Kolodkin AL, Ghosh A. Patterning of cortical efferent projections by semaphorin-neuropilin interactions. Science (80-). 1998;282(5395):1904–1906. doi:10.1126/science.282.5395.1904

51. Gao PP, Yue Y, Zhang JH, Cerretti DP, Levitt P, Zhou R. Regulation of thalamic neurite outgrowth by the Eph ligand ephrin-A5: implications in the development of thalamocortical projections. Proc Natl Acad Sci U S A. 1998;95(9):5329–5334. doi:10.1073/pnas.95.9.5329

52. Baudet M-L, Zivraj KH, Abreu-Goodger C, et al. miR-124 acts through CoREST to control onset of Sema3A sensitivity in navigating retinal growth cones. Nat Neurosci. 2012;15(1):29–38. doi:10.1038/nn.2979

53. Shewan D, Dwivedy A, Anderson R, Holt CE. Age-related changes underlie switch in netrin-1 responsiveness as growth cones advance along visual pathway. Nat Neurosci. 2002;5(10):955–962. doi:10.1038/nn919

54. Skaliora I, Singer W, Betz H, Püschel AW. Differential patterns of semaphorin expression in the developing rat brain. Eur J Neurosci. 1998;10(4):1215–1229. doi:10.1046/j.1460-9568.1998.00128.x

55. Sturrock RR. Myelination of the mouse corpus callosum. Neuropathol Appl Neurobiol. 1980;6(6):415–420. doi:10.1111/j.1365-2990.1980.tb00219.x

56. Goebbels S, Wieser GL, Pieper A, et al. A neuronal PI(3,4,5)P3-dependent program of oligodendrocyte precursor recruitment and myelination. Nat Neurosci. 20(1):10–15. doi:10.1038/nn.4425

57. Tomassy GS, Berger DR, Chen H-HH, et al. Distinct Profiles of Myelin Distribution Along Single Axons of Pyramidal Neurons in the Neocortex. Science (80-). 2014;344(6181):319–324. doi:10.1126/science.1249766

58. Micheva KD, Wolman D, Mensh BD, et al. A large fraction of neocortical myelin ensheathes axons of local inhibitory neurons. Elife. 2016;5. doi:10.7554/eLife.15784

59. Ishii K, Kubo KI, Endo T, et al. Neuronal heterotopias affect the activities of distant brain areas and lead to behavioral deficits. J Neurosci. 2015. doi:10.1523/JNEUROSCI.3648-14.2015

60. Roper SN, Gilmore RL, Houser CR. Experimentally induced disorders of neuronal migration produce an increased propensity for electrographic seizures in rats. Epilepsy Res. 1995;21(3):205–219. doi:10.1016/0920-1211(95)00027-8

61. Nieto M, Monuki ES, Tang H, et al. Expression of Cux-1 and Cux-2 in the subventricular zone and upper layers II-IV of the cerebral cortex. J Comp Neurol. 2004;479(2):168–180. doi:10.1002/cne.20322

62. Chmielowska J, Stewart MG, Bourne RC. gamma-Aminobutyric acid (GABA) immunoreactivity in mouse and rat first somatosensory (SI) cortex: description and comparison. Brain Res. 1988;439(1-2):155–168. doi:10.1016/0006-8993(88)91472-2

63. Meyer HS, Schwarz D, Wimmer VC, et al. Inhibitory interneurons in a cortical column form hot zones of inhibition in layers 2 and 5A. Proc Natl Acad Sci. 2011;108(40):16807–16812. doi:10.1073/pnas.1113648108

64. Gabel LA, LoTurco JJ. Layer I Ectopias and Increased Excitability in Murine Neocortex. J Neurophysiol. 2002;87(5):2471–2479. doi:10.1152/jn.2002.87.5.2471

65. Manent JB, Wang Y, Chang Y, Paramasivam M, LoTurco JJ. Dcx reexpression reduces subcortical band heterotopia and seizure threshold in an animal model of neuronal migration disorder. Nat Med. 2009;15(1):84–90. doi:10.1038/nm.1897

66. Chevassus-Au-Louis N, Rafiki A, Jorquera I, Ben-Ari Y, Represa A. Neocortex in the hippocampus: An anatomical and functional study of CA1 heterotopias after prenatal treatment with methylazoxymethanol in rats. J Comp Neurol. 1998;394(4):520–536. doi:10.1002/(SICI)1096-9861(19980518)394:4<520::AID-CNE9>3.0.CO;2-3

67. Jenner AR, Galaburda AM, Sherman GF. Connectivity of ectopic neurons in the molecular layer of the somatosensory cortex in autoimmune mice. 2000;10(10):1005–1013.

68. Chevassus-Au-Louis N, Congar P, Represa A, Ben-Ari Y, Gaïarsa JL. Neuronal migration disorders: heterotopic neocortical neurons in CA1 provide a bridge between the hippocampus and the neocortex. Proc Natl Acad Sci U S A. 1998;95(17):10263–10268. doi:10.1073/PNAS.95.17.10263

69. Schottler F, Fabiato H, Leland JM, et al. Normotopic and heterotopic cortical representations of mystacial vibrissae in rats with subcortical band heterotopia. Neuroscience. 2001. doi:10.1016/S0306-4522(01)00395-5

70. Romero DM, Bahi-Buisson N, Francis F. Genetics and mechanisms leading to human cortical malformations. Semin Cell Dev Biol. 2018;76:33–75. doi:10.1016/J.SEMCDB.2017.09.031

71. Saito T. In vivo electroporation in the embryonic mouse central nervous system. Nat Protoc. 2006;1(3):1552–1558. doi:10.1038/nprot.2006.276

72. Kim J-Y, Ash RT, Ceballos-Diaz C, et al. Viral transduction of the neonatal brain delivers controllable genetic mosaicism for visualising and manipulating neuronal circuits in vivo. Eur J Neurosci. 2013;37(8):1203–1220. doi:10.1111/ejn.12126

73. Phifer CB, Terry LM. Use of hypothermia for general anesthesia in preweanling rodents. Physiol Behav. 1986;38(6):887–890.

74. Patel TP, Man K, Firestein BL, Meaney DF. Automated quantification of neuronal networks and single-cell calcium dynamics using calcium imaging. J Neurosci Methods. 2015;243:26–38. doi:10.1016/j.jneumeth.2015.01.020

75. Li X, Ouyang G, Usami A, Ikegaya Y, Sik A. Scale-Free Topology of the CA3 Hippocampal Network: A Novel Method to Analyze Functional Neuronal Assemblies. Biophys J. 2010;98(9):1733–1741. doi:10.1016/J.BPJ.2010.01.013

76. Li X, Cui D, Jiruska P, Fox JE, Yao X, Jefferys JGR. Synchronization Measurement of Multiple Neuronal Populations. J Neurophysiol. 2007;98(6):3341–3348. doi:10.1152/jn.00977.2007

